# Effects of interspecies interactions on marine community ecosystem function

**DOI:** 10.1101/2022.08.26.505414

**Authors:** Michael Daniels, Astrid K.M. Stubbusch, Noelle A. Held, Olga T. Schubert, Martin Ackermann

## Abstract

Microbial communities perform key ecosystem processes collectively. One such process is the degradation of carbohydrate polymers, which are the dominant pool of organic carbon in natural environments. Carbohydrate polymers are often degraded in a stepwise manner. Individual steps are performed by different microbial species, which form trophic cascades with carbon polymers at the bottom and fully oxidised carbon at the top. It is widely believed that these trophic cascades are hierarchically organised, where organisms at each level rely on organisms at the levels below. However, whether and how the higher-level organisms can also affect processes at the lower levels is not well understood. Here we studied how carbohydrate polymer degradation mediated by secreted enzymes is affected by species at higher trophic levels, i.e., species that cannot produce the enzymes for polymer degradation but can grow in presence of the polymer degraders. We used growth and enzyme assays in combination with transcriptomics to study how chitin degradation by a number of Vibrio strains is affected by the presence of different cross-feeders that consume metabolic by-products. We found that interactions between the degraders and cross-feeders influence the rate of chitin degradation by the community. Furthermore, we show that this is a result of changes in chitinase expression by degraders. Overall, our results demonstrate that interactions between species can influence key ecosystem functions performed by individuals within microbial communities. These results challenge the perspective that trophic cascades based on metabolically coupled microbial communities are unidirectional and provide mechanistic insights into these downstream interactions.

## Introduction

In natural environments microorganisms form multispecies communities that are important contributors to a wide range of ecosystem functions (1). A central question is how these ecosystem functions depend on the composition of a community and its diversity (2). It has been found that diversity is a key component for the resilience and general productivity of microbial communities. While some community level properties such as total biomass production generally increase to a certain extent with the total number of species and therefore overall diversity (3, 4, 5, 6), other studies have found that certain functions are carried out only by a subset of species in a community (7, 8). For example, studies on diversity in aquatic microbial communities have shown that the metabolic function of polysaccharide degradation is controlled by single clades and their metabolic traits rather than by species richness of the total community (9).

In nature, biopolymers are important nutrient sources for microbial communities. The most abundant biopolymers in terrestrial systems are the polysaccharides cellulose (10) and lignin (11), whereas in aquatic systems the most abundant biopolymer is the aminopolysaccharide chitin (12). These polymeric substances are synthesised by multicellular organisms and form structural components of a plethora of animals and plants. Once the producing organism dies, these polymers become available for microbial communities to be consumed as carbon source whereby they are eventually remineralised back to inorganic carbon dioxide. Thereby, the collective metabolism of microbial communities that degrade polymers plays a vital role in the global cycling of nutrient elements.

Many natural polymers are carbohydrate chains of repeating subunits mostly linked by beta-1,4 glycosidic bonds. These polymers have high molecular weights, therefore they require cleavage by extracellular enzymes prior to uptake by microbes (13). While these enzymes are produced by specialised microbes, the resulting mono- and dimeric subunits released from the polymers can be taken up and metabolized by many microbial organisms. Hence, the extracellular degradation of polymers generates a pool of nutrients in the local environment that is also accessible for microbes that lack the necessary enzymes for polymer degradation. These non-degraders fall into two groups, consumers and cross-feeders. Consumers can take up the primary degradation products, i.e., the mono- or dimers, and hence compete with the degraders for these degradation products. Cross-feeders are unable to consume the primary degradation products, instead they rely on metabolic by-products such as acetate or amino acids that are excreted by the degraders and consumers. Therefore, the polymer degraders - via their initial release of enzymes - form the foundation of a multi-level trophic cascade.

In the marine environment the most abundant polysaccharide is chitin. Chitin is composed of N-acetylglucosamine subunits and builds the structural component of the exoskeletons of crustaceans. Through the natural life cycles of these organisms’ chitin exoskeletons eventually decay into particles and constitute an important nutrient source for marine microbial species. Polysaccharide sources like marine particles are usually colonized by a diverse community of microorganisms that assemble on the particle surface and can be metabolically tightly coupled. Recent work showed that, typically, food chains structured around polymer degradation are hierarchically organised (14). In these trophic cascades, non-degrading species benefit from initial chitin breakdown by degraders, which express a wide array of chitinolytic enzymes.

Bacterial chitinolytic enzymes (hereafter called chitinases) are grouped into glycosyl hydrolase families GH18, GH19 and GH20 based on their amino acid sequence (15). Furthermore, these bacterial chitinases are grouped into two broad subclasses: endo- and exo-chitinases. Endochitinases cleave off smaller oligomers (16). These oligomers cannot be readily taken up by microbial cells and need to be further broken down into dimers or monomers that can be imported and metabolised (17). Exochitinases cleave off monomers or dimers and can be further subdivided into chitobiosidases (18) and β-1,4-*N*-acetyglucosaminidases (19). Chitobiosidases produce the chitin dimer di-acetylchitobiose (chitobiose) while β-1,4-*N*-acetyglucosaminidases (hereafter N-acetyglucosaminidases) produce the monomer *N*-acetylglucosamine (GlcNAc).

Here, we investigate whether and how species without the ability to degrade a polymer influence polymer degradation on a community level. Specifically, we focus on the impact of non-chitinolytic species on the hydrolytic activity of chitin degraders. The underlying question we aim to answer is whether bacterial species without the genetic repertoire to perform a given function can influence that function when measured at the level of consortia. We show that species-specific interactions between degraders and non-degraders lead to increased chitinase activity in co-cultures. Using a transcriptomics approach, we further show that in co-cultures, gene expression profiles in degrader species are specific to the cross-feeder species, with higher expression of chitinases in the presence of some but not other cross-feeders. Overall, our study provides novel insights on how specific interspecies interactions influence community level function on the model of chitin remineralization.

## Results and Discussion

### Chitin degraders produce a diverse repertoire of chitinolytic enzymes

To study the roles of and interactions between members of chitin-degrading microbial communities, we first focused on the community members at the bottom of the trophic hierarchy, the degraders. We first investigated the activity of different chitinases that are produced as degraders grow on chitin as a sole carbon source. To this end, we selected ten closely related chitin degraders belonging to the family of Vibrionaceae and measured the activity of chitinases in their supernatants using enzymatic hydrolysis substrates that contain 4-methylumbelliferone (Methods). The resulting products indicate the activity of three different chitinase groups, those that produce the monomer *N*-acetylglucosamine (GlcNAc), those that produce the dimer chitobiose, and those that produce chitin oligomers.

We found that for all degrader species except *V. fisherii* ZF211 the highest enzymatic activity was displayed for chitobiosidases, the enzymes that cleave dimers off chitin (Figure 1). The activity of the other two enzyme classes was generally lower, again with the exception of *V. fischeri* ZF211, which showed comparable activity of chitobiosidases and *N*-acetyglucosaminidases. Taken together, these findings suggest that in monocultures of the Vibrionaceae strains tested here, the dimer chitobiose, rather than the monomer GlcNAc, is the dominant chitin degradation product.

**Figure 1:**
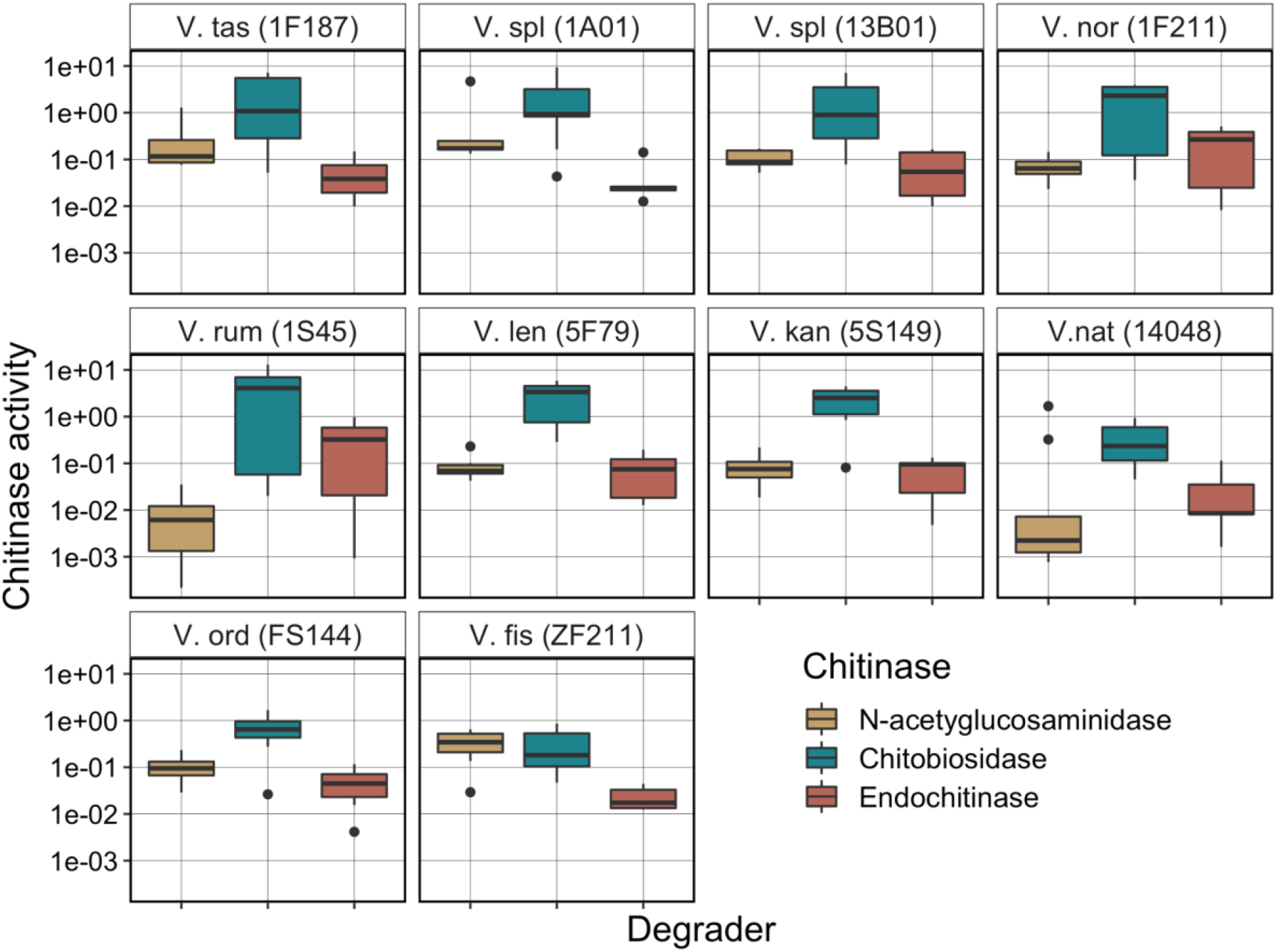
Enzymatic activities of the three general chitinase groups vary among degrader species. For each of 10 chitin degraders (monocultures), the enzymatic activities of β-1,4–*N*-acetyglucosaminidases (yellow), chitobiosidases (green) and endochitinases (red) are shown. Chitinase activity reflects enzyme units per microliter per growth yield (OD) of culture.

These results are relevant in light of previous findings, showing that chitobiose is a potent chemoattractant (20, 21) and that different degradation products lead to specific gene-expression responses in *Vibrio* (22, 23).

### Chitinase activity, growth rate, and yield are tightly coupled

In microbial systems that rely on polymer degradation for nutrient availability, growth and enzyme production are tightly coupled (24, 25). Increases in hydrolytic enzyme production and activity were found to lead to increased cell growth; furthermore, the number of degrading cells tends to be positively correlated with the amount of enzymes produced. In order to understand the relationship between enzyme activity, growth rate and yield among chitin degrading species, we quantified the correlation between enzyme activity and growth dynamics in monocultures of our ten selected degraders.

We found that the activity of the three chitinase groups are differently correlated with growth (Figure 2). Chitobiosidases showed the highest correlation with yield in our bacterial populations, while *N*-acetyglucosaminidases had low correlations for both yield and growth rate. Furthermore, the degrader monocultures displayed a high degree of correlation between growth rate and yield (Figure S1). These findings show that chitinase activity and population level growth in degrader monocultures are tightly and positively coupled. The species with highest enzyme activity display the most growth, but the exact causal relationship and the mechanisms underlying these correlations are still unknown. For chitobiosidases and endochitinases the positive correlation with yield and growth rate persisted even when enzyme activity was normalised by total yield (as a proxy for biomass), establishing that strains that produced higher levels of these chitinases *per cell* achieved faster growth and larger population sizes (Figure 2C and 2D). This indicates a general positive association between these enzymes and growth parameters of these strains.

**Figure 2:**
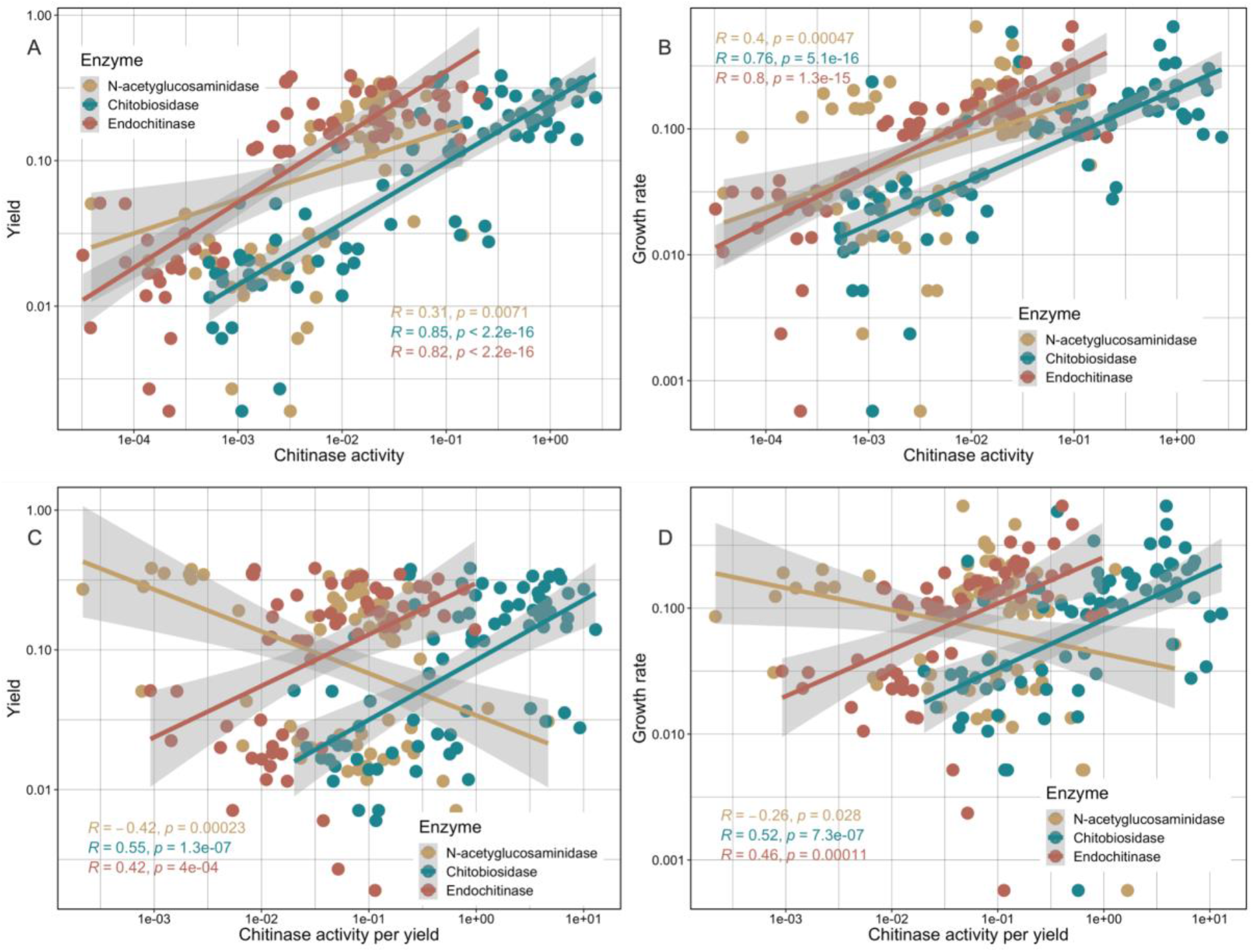
Growth dynamics and catalytic activity are positively correlated. Chitinase activity is positively correlated with yield and growth rates of degrader monocultures. Pearson’s correlation and 95% CI are shown. Activity of individual enzyme classes as a function of degrader monoculture yield (A) or growth rate (B) and activity of individual enzyme classes normalized by yield as a function of degrader monoculture yield (C) or growth rate (D). β− 1,4-*N*-acetyglucosaminidases (yellow), chitobiosidases (green) and endochitinases (red).

### Non-degrading cross-feeders selectively influence enzyme activity on the community level

In natural polymer-degrading communities, the process of polymer degradation is performed by a small subset of microbial species (26). It is generally assumed that non-degraders do not contribute to overall enzymatic activity at the community level (27). However, as described above, degrader growth rate and yield correlated with chitinase activity (Figure 2). To investigate whether the presence of individual non-degrading species changes a community’s properties, we co-inoculated a variety of different degraders with non-degrading cross-feeders. We found that non-degraders influenced growth rate and yield on a community level (Figure S2 & S3). This raises the question whether the presence of cross-feeders can also affect chitinase activity on the community level. In order to address this question, we investigated the impact of non-degrading species on the enzymatic activity of the degraders.

In 10 out of 53 degrader-cross-feeder pairs the presence of the cross-feeder led to statistically significant increases or decreases in total chitobiosidase activity (Figure 3A). We found that certain cross-feeders, particularly the two *Alteromonas* strains tested, increased chitobiosidase activity of several of the degraders (Figure 3A and 3B). In contrast, for example, the presence of *Neptunomonas* reduced chitinase activity in almost half of the co-cultures tested or had no significant effect (Figure 3A and 3B). N-acetylglucosaminidase activity (Figure S4) and endochitinase (Figure S5) activities were also found to be different in certain co-cultures. Endochitinase activity showed comparable patterns to chitobiosidase activity. There, we found eleven pairs where the presence of cross-feeders leads to statistically significant changes, with the two *Alteromonas* strains increasing endochitinase activity in *V. natriegens* and *V. splendidus* (Figure S5). N-acetylglucosaminidase activity, in contrast, was decreased in all 14 cases where we found statistically significant changes in the enzyme activity (Figure S4). There, the presence of the two *Alteromonas* strains (for which we found increases in the other two chitinase groups) lead to decreases in enzyme activity. Notably, in co-cultures of *V. fisherii* ZF211 and *V. splendidus* 13B01 we found increases in one chitinase group and decreases in another. Furthermore, we found that the presence of cross-feeders can influence community level yield (Figure S2) and growth rate (Figure S3). Among the cross-feeders with the strongest effect on community level yield were *Alteromonas* and *Neptunomonas*.

**Figure 3:**
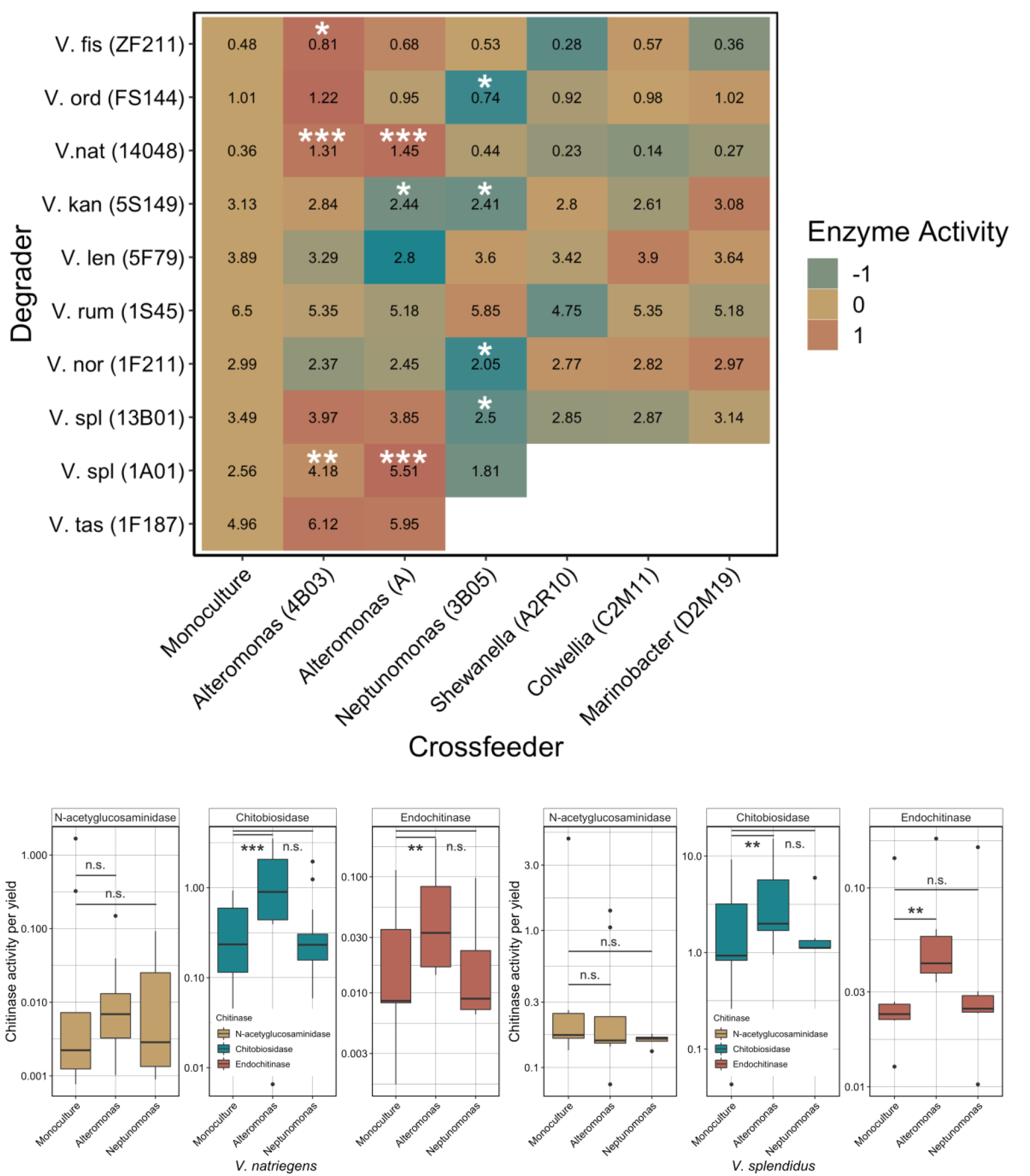
Crossfeeders selectively influence chitinase activity of communities. A) Heatmap of enzymatic activity per OD for various degraders (vertical-axis) in monoculture (first column) or in co-culture with different cross-feeders (horizontal-axis). Numbers indicate total chitinase activity per total OD per µL (Methods). Colours represent activities in co-cultures relative to the degrader monoculture. Yellow colours indicate little to no change compared to the degrader monoculture. Green colours represent a reduced enzymatic activity. Red colours represent an increase in enzyme activity. White panels indicate combinations that were not measured. Stars represent significant changes in growth between the monoculture and respective co-culture: * = *p* < 0.05, ** = *p* < 0.01, *** = *p* < 0.001 (ANOVA). Details about the underlying statistical analysis can be found in the supplementary material (Table S3). B) Enzymatic activity of *N-*acetylglucosaminidases (yellow), chitobiosidases (green) and endochitinases (red) per yield of *V. natriegens* monocultures (left) compared to co-cultures with *Alteromonas* (middle) and *Neptunomonas* (right). C) Enzymatic activity for *V. splendidus* monocultures (left) of *N-*acetylglucosaminidases (yellow), chitobiosidases (green) and endochitinases (red) per yield compared to co-cultures with *Alteromonas* (middle) and *Neptunomonas* (right).

### Species-specific interactions influence community level processes

We next set out to explore by which mechanism the presence of certain cross-feeding species induces chitinase activity in the communities. We hypothesized that the removal of excreted metabolites by cross-feeders could directly affect the catalytic activity of individual chitinases secreted by the degraders through the release of product inhibition. Alternatively, the presence of the cross-feeder could induce gene expression changes in the degrader, e.g., leading to increased expression or secretion of chitinases. To study this, we used degrader-cross-feeder co-cultures involving *Neptunomonas* and *Alteromonas* strains. The individual interactions between *Neptunomonas* and *V. splendidus* and *Alteromonas* and *V. natriegens* strains have previously been studied in detail (28, 29). These studies revealed that both the *Alteromonas* and *Neptunomonas* strain were cross-feeders in the sense that they are unable to degrade chitin or consume primary degradation products such as mono- or dimers and instead rely on metabolic by-products excreted by the degraders, such as acetate for carbon sources.

To understand the mechanism by which these two cross-feeders affect chitinase activity, we compared global gene expression profiles of two different degraders in monoculture (*V. splendidus* 1A01 and *V. natriegens* ATCC 14048) with their gene expression profiles in co-culture with one of two cross-feeders (*Alteromonas* 4B03 and *Neptunomonas* 3B05) (Figure 4). We found marked effects of the presence of cross-feeders on gene expression of the degraders. The transcriptomes of *V. natriegens* showed that the expression of 1,499 genes (32% out of 4,640 quantified genes) was significantly different (*p* < 0.05) in the presence of a cross-feeder compared to the monoculture (Figure 4A). For *V. splendidus*, we found 1,965 genes (38% out of 5,237 quantified genes) to be differentially expressed in the presence of the cross-feeder (Figure 4B). The gene expression differences between the co-cultures with the cross-feeder species were overall similar but with 172 genes and five genes differentially regulated in *V. natriegens* and *V. splendidus*, respectively (Figure 4).

**Figure 4:**
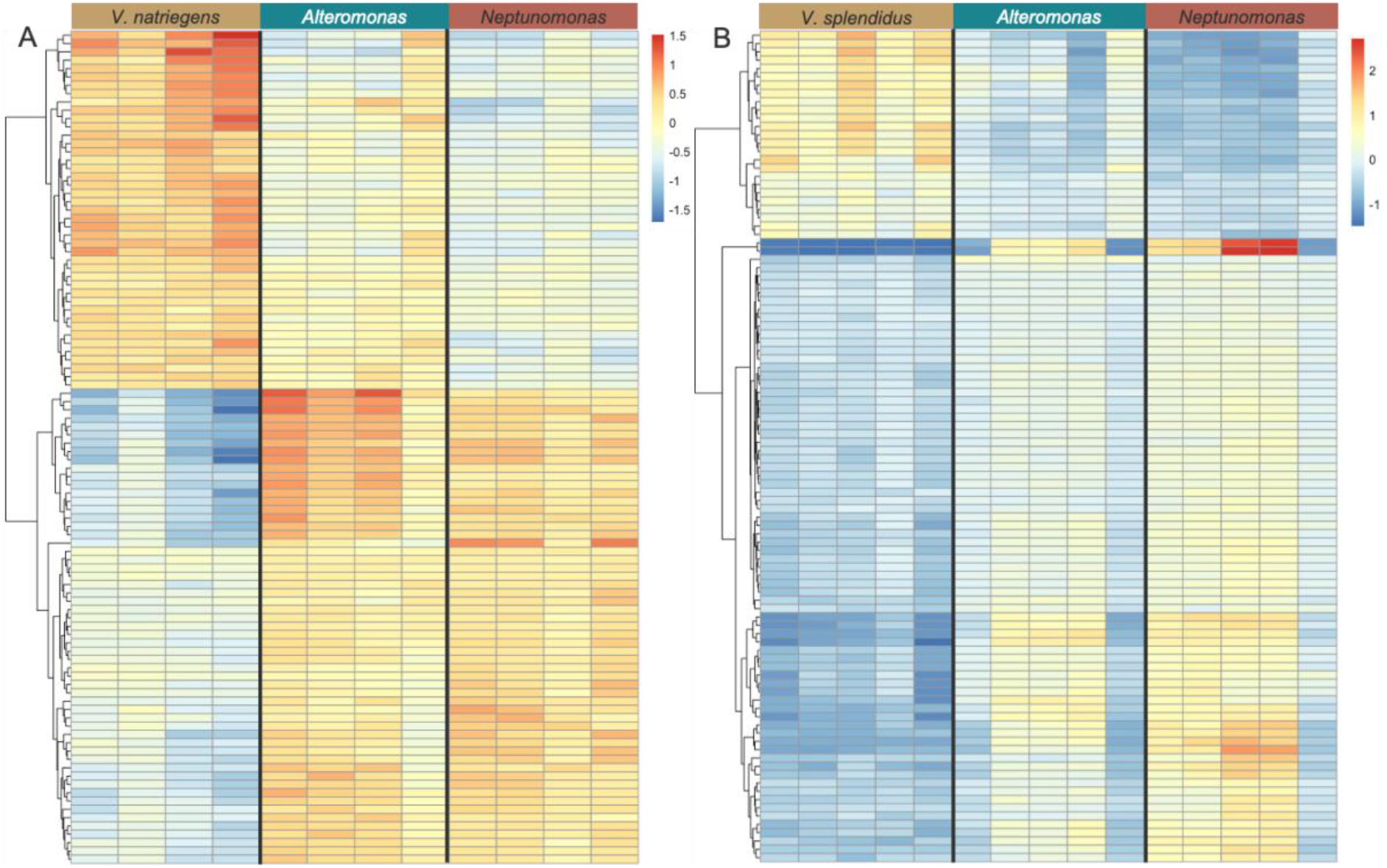
The presence of cross-feeders changes gene expression profiles in degraders. Degraders show large differences in gene expression profiles in co-cultures with cross-feeders compared to monocultures. Colors indicate log2 fold change (DESeq2) A) Top 100 most significantly differentially expressed genes (regularized logarithm (rlog) function, DESeq2, sorted by adjusted *p*-value) for *V. natriegens* monocultures, compared to a co-culture with *Alteromonas* or *Neptunomonas*. In total 1,107 out of 4,641 genes were found to be differentially expressed between the mono- and the co-culture with *Alteromonas*, while 1,585 out of 4,641 genes were differentially expressed in the presence of *Neptunomonas* compared to the monoculture (*p* < 0.05). B) Top 100 most significantly differentially expressed genes (Regularized logarithm (rlog) function, DESeq2, adjusted *p* value) for *V. splendidus* monocultures, compared to a co-culture with *Alteromonas* and *Neptunomonas*. In total 690 out of 5,237 genes were found to be differentially expressed in between the mono- and the co-culture with *Alteromonas*, while 1,869 out of 5,237 genes were differentially expressed in the presence of *Neptunomonas* compared to the monoculture (*p* < 0.05).

### Increases expression of chitinase genes in *Vibrio natriegens* in the presence of *Alteromonas*

We found that in the presence of *Alteromonas* three of the seven chitinases of the degrader *V. natriegens* were upregulated compared to the degrader monoculture (Figure 5A). This is consistent with the finding that *Alteromonas* increases community level chitinase activity (Figure 3). Furthermore, we found that various other enzymes important for polymer degradation were upregulated, including glycosyl hydrolases (Figure S7), a general enzyme family that catalyses the hydrolysis of complex carbohydrates (30) as well as auxiliary enzymes (Figure S8) like lytic polysaccharide mono-oxigenases that aid in the degradation of polymers through interaction with degrading enzymes (31). Additionally, we found that not only the chitinases but chitin catabolism more generally (KEGG: “amino sugar and nucleotide sugar pathway”) was upregulated in *V. natriegens* (Figure S6) when grown in the presence of *Alteromonas*. In contrast, we found that the presence of *Neptunomonas* does not consistently induce chitinase expression in the degrader (Figure 5B), consistent with our finding that *Neptunomonas* does not increase chitinase activity in co-culture. Taken together our findings show that the gene expression response of *V. natriegens* is specific to the individual cross-feeding species, leading to specific differences in overall chitin degradation activity at the community level.

**Figure 5:**
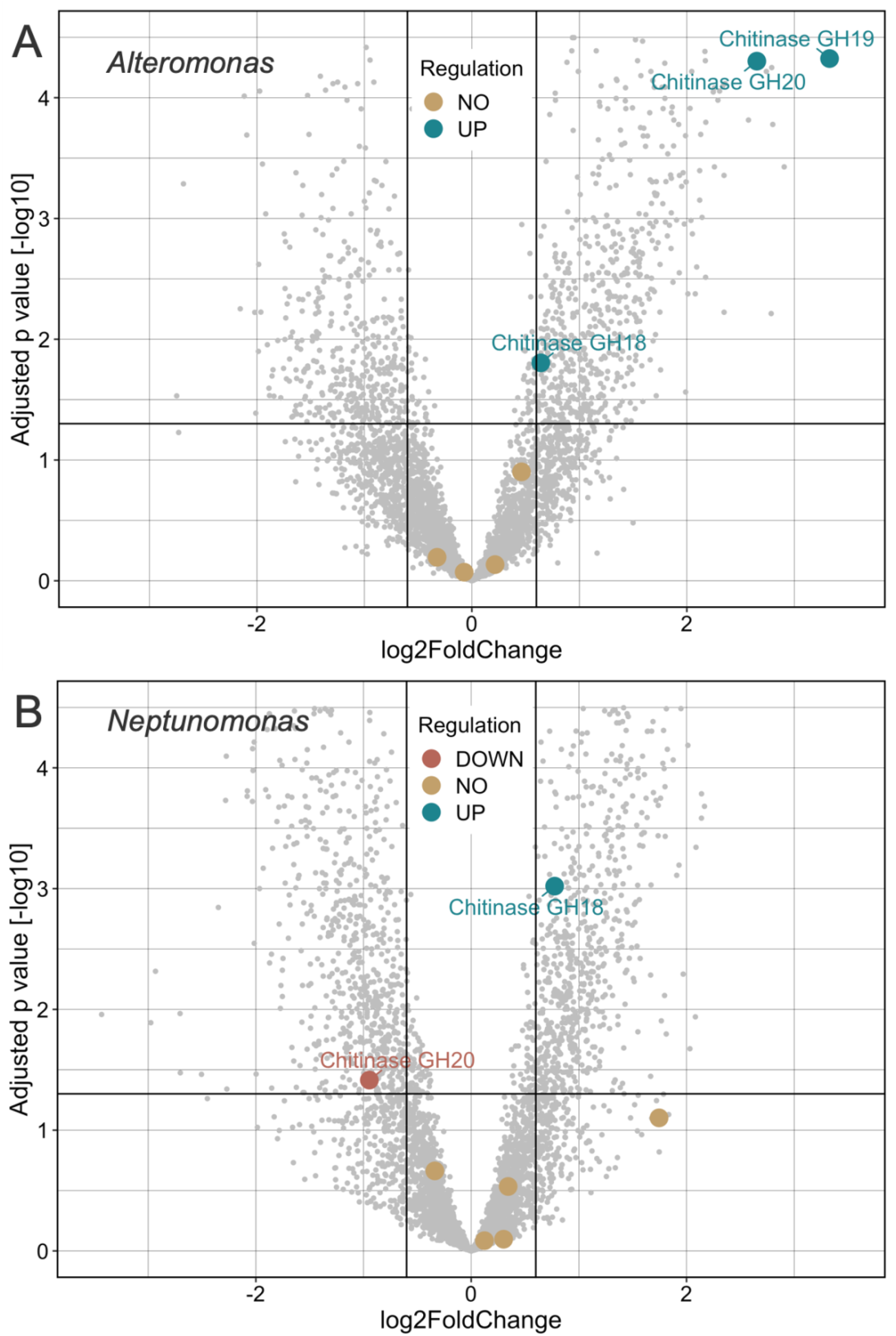
Differential expression of chitinases in *V. natriegens*. A) *Alteromonas* induces expression of several chitinases in *V. natriegens*. Coloured dots indicate individual chitinases found in the genome of *V. natriegens*, black lines indicate significance cut offs (vertical lines indicate |log2 fold change| = 0.6, the horizontal line indicates *p* < 0.05) colours indicate differential expression based on the cut-off criteria; down-regulation (red), no difference (yellow), up-regulation (green). Three out of seven chitinases were significantly upregulated in the presence of *Alteromonas*. B) *Neptunomonas* does not consistently induce chitinase expression in *V. natriegens*. Differences in chitinase expression in *V. natriegens* in the presence vs absence of *Neptunomonas*. Coloured dots indicate individual chitinases found in the genome of *V. natriegens*, black lines indicate significance cut-offs (vertical lines indicate |log2 fold change| = 0.6, the horizontal line indicates *p* < 0.05) colours indicate differential expression based on the cut-off criteria; down-regulation (red), no difference (yellow), up-regulation (green). Two out of seven chitinases were differentially expressed in the presence of *Neptunomonas*. One was up regulated and one was down regulated.

### Cross-feeders influence degraders’ expression of functional gene groups

Polymer degradation is commonly linked with collectivity at the level of degrader populations. We recently found that when cells collectively degrade polymers they show a low degree of motility (32). This indicates that polymer degradation is a cell density dependent function where cells engage in the production of public goods and benefit from microcolony formation (33). In *Vibrio* species, group level behaviour is associated with the expression of quorum sensing gene clusters (34) that regulate various density dependent functions such as virulence, biofilm formation and polymer degradation. Indeed, we found that in the presence of *Alteromonas, V. natriegens* displayed a significant increase in expression of genes associated with general motility (Figure S9). We also found a slight decrease in the expression of genes associated with quorum sensing (Figure S10). Furthermore, some of the genes in the quorum sensing pathway that were upregulated in the presence of *Alteromonas*, such as LuxO, are known repressors of high-density dependent processes in *Vibrio* species (35). This further hints towards a general downregulation of quorum sensing dependent functions for *V. natriegens* in the presence of *Alteromonas*. In order to test this hypothesis, further studies are necessary.

### Interactions between *Vibrio splendidus* and cross-feeders

So far, we have shown that the effect of the cross-feeder on a degrader is species-specific, both on the level of growth rate and yield as well as on the level of gene expression. Next, we set out to assess the generalizability of the relationship by asking whether the same cross-feeders have similar effects on different degraders. To this end, we assessed the differences in transcriptional responses in *V. splendidus* in isolation compared to a co-culture with one of the two cross-feeders. When measuring chitinase activity of these pairs, we found comparable dynamics to the degrader *V. natriegens* (Figure 3). There the presence of *Alteromonas* led to increased chitinase activity while the presence of Neptunomonas had no significant effect (Figure 3). Therefore, we first investigated the changes in chitinase expression in the presence of these cross-feeders.

This time, we tested the effect of the cross-feeders *Alteromonas* and *Neptunomonas* on the chitin degrader *V. splendidus*. We found that *Alteromonas* leads to higher expression of chitinase genes in *V. splendidus*, consistent with our finding that *Alteromonas* also increases the community level enzyme activity in co-culture with *V. splendidus* (Figure 6A). Contrary to our hypothesis, we found that *Neptunomonas* also leads to higher expression of various chitinases in *V. splendidus* (Figure 6B). This was surprising, because our earlier experiments had shown decreased rather than increased chitinase activity on the community level in the presence of *Neptunomonas* (Figure 2).

**Figure 6:**
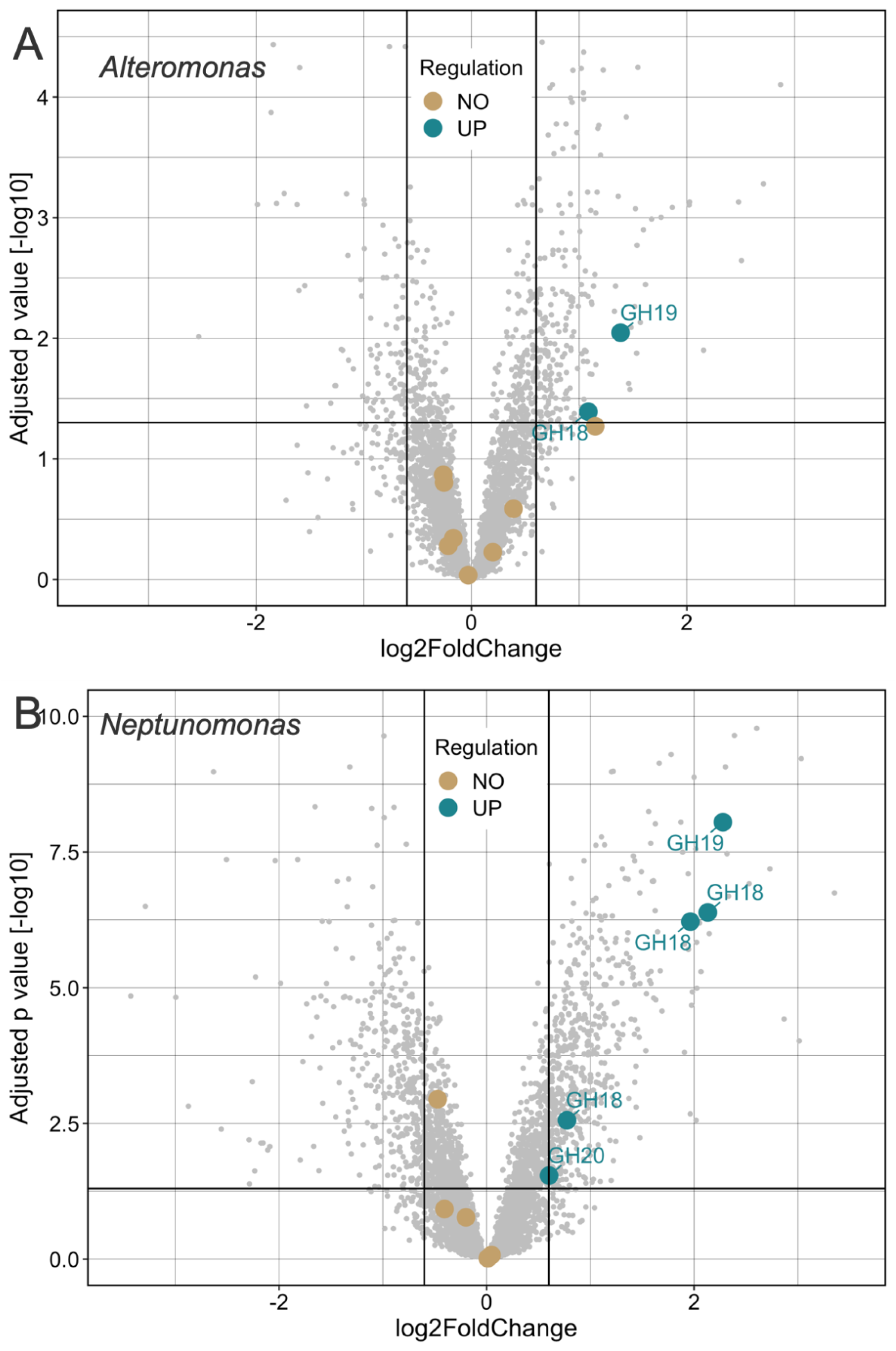
Differential expression of chitinases in *V. splendidus*. A) *Alteromonas* induces expression of chitinases in *V. splendidus*. Differences in chitinase expression in *V. splendidus* in the presence vs absence of *Alteromonas*. Dots indicate individual chitinases found in the genome of *V. splendidus*, black lines indicate significance cut offs (vertical lines indicate |log2 Fold Change| = 0.6, the horizontal line indicates *p* < 0.05) colours indicate differential expression based on the cut off criteria; down-regulation (red), no difference (yellow), up-regulation (green). Two out of ten chitinases were significantly upregulated in the presence of *Alteromonas*. B) *Neptunomonas* induces expression of chitinases in *V. splendidus*. Differences in chitinase expression in *V. splendidus* in the presence vs absence of *Neptunomonas*. Dots indicate individual chitinases found in the genome of *V. splendidus*, black lines indicate significance cut offs (vertical lines indicate |log2 Fold Change| = 0.6, the horizontal line indicates *p* < 0.05) colours indicate differential expression based on the cut off criteria; down-regulation (red), no difference (yellow), up-regulation (green). Five out of ten chitinases were significantly upregulated in the presence of *Neptunomonas*.

### Secreted chitinases can serve as nutrient sources

Our findings above indicate that the relationship between species interactions, chitinase expression and enzyme activity is not trivial and suggest that the influence of cross-feeders on community-level chitinase activity goes beyond affecting chitinase gene expression in the degrader species. The abundance of extracellular chitinases is determined not just by their production and secretion but also by their stability. Indeed, chitinases themselves may be subject to degradation, e.g., by proteases secreted by community members in order to release amino acids that can in turn be consumed as nutrient sources. To test whether chitinases can themselves be used as a nutrient source, we grew our four focal strains in media containing lyophilised enzyme mixes as sole carbon source. We found that the four strains can grow on these enzymes to various degrees (Figure S11).

Next, we measured how the activity of N-acetylglucosaminidases, chitobiosidases, and endochitinases in the media changes when used as sole carbon source to grow our communities. We found that a co-culture with *V. natriegens* and *Alteromonas* decreased the activity of N-acetylglucosaminidases compared to the degrader monoculture when grown on enzymes as a substrate (Figure 7A). This result was consistent with our findings that *Alteromonas* reduced the activity of N-acetylglucosaminidases in many co-cultures that grow on chitin (Figure S5). Also, in co-cultures with *V. splendidus* we found a decrease in enzyme activity for N-acetylglucosaminidases of the substrate in the presence of *Alteromonas* compared to a monoculture (Figure 7B). The selective decrease in activity for N-acetylglucosaminidases (Figure 7A & 7B) in co-cultures with *Alteromonas* indicates a preference for certain specific enzymes as substrates. The induction of chitobiosidase expression (Figure 5A & 6A) combined with the selective degradation of N-acetylglucosaminidases could explain our findings that in *V. fischeri* (ZF211), we simultaneously observe increases for chitobiosidase activity (Figure 2) and decreases for N-acetylglucosaminidase activity (Figure S5) in co-cultures with *Alteromonas*. For co-cultures with *Neptunomonas* we found no change in the activity of N-acetylglucosaminidases but a decrease for chitobiosidase activity of the substrate (Figure 7B). This result is consistent with our findings that *Neptunomonas* reduced the activity of N-acetylglucosaminidases in many co-cultures that grow on chitin (Figure 2).

**Figure 7:**
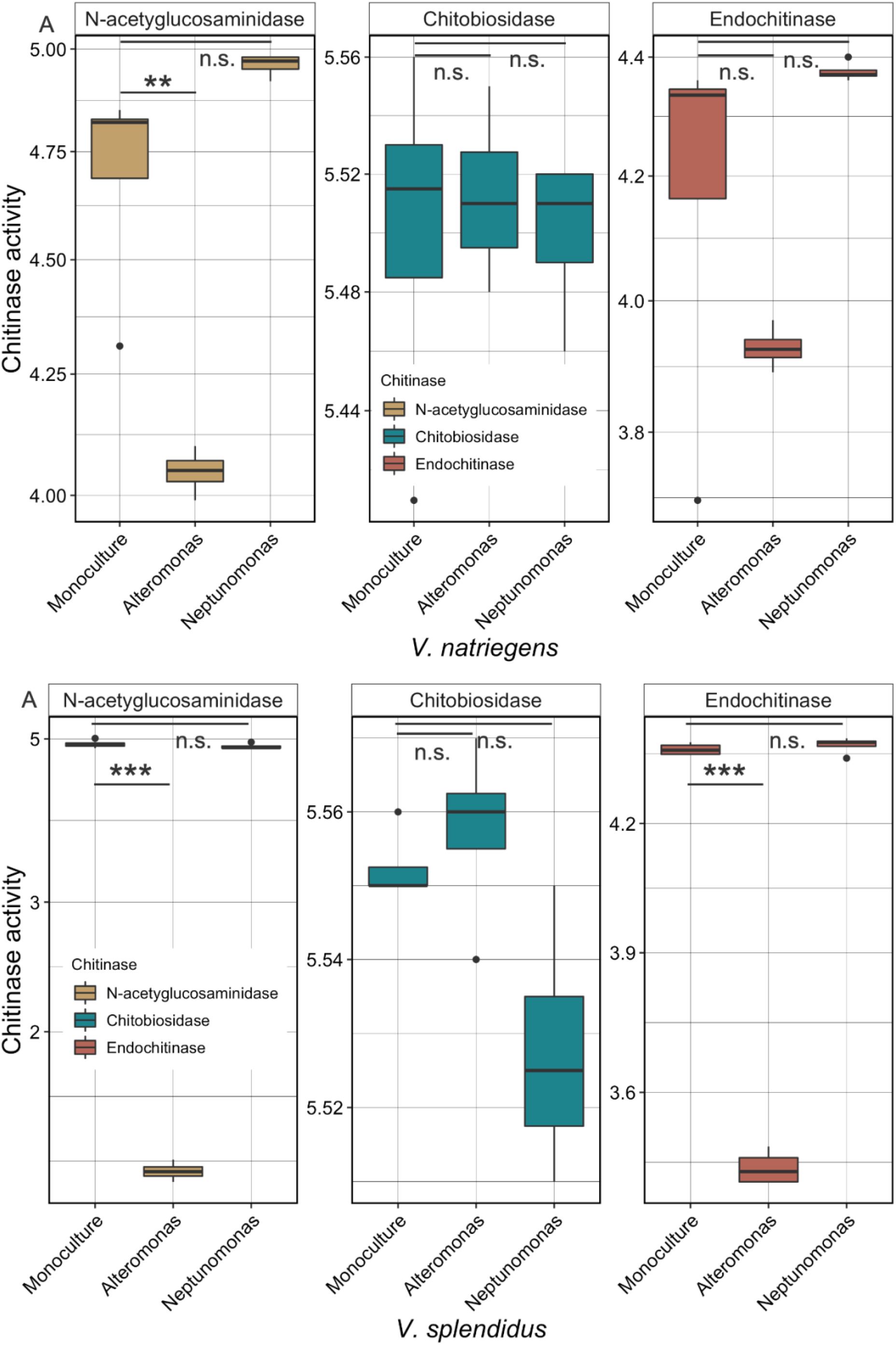
Communities consume enzymes selectively. Enzymatic activity of *N*-acetyglucosaminidases (yellow), chitobiosidases (green), and endochitinases (red) changes depending on community composition. (A) Compared to *V. natriegens* monocultures (left) the co-culture with *Alteromonas* (middle) leads to a decrease in *N*-acetyglucosaminidase activity of the substrate. The presence of *Neptunomonas* (right) has no effect. (Welch Two Sample t-test; N = 4; *N*-acetyglucosaminidase: mean of monoculture = 4.70 U/µL, mean of co-culture *Alteromonas* = 4.05 U/µL; *t* = 4.94, *p* value = 0.01, mean of co-culture *Neptunomonas* = 4.96 U/µL; *t* = -1.99, *p* value = 0.14. Chitobiosidases: mean of monoculture = 5.50 U/µL, mean of co-culture *Alteromonas* = 5.51 U/µL; *t* = - 0.36, *p* value = 0.74, mean of co-culture *Neptunomonas* = 5.50 U/µL; *t* = 0, *p* value = 1.00. Endochitinases: mean of monoculture = 4.18 U/µL, mean of co-culture *Alteromonas* = 3.93 U/µL; *t* = 1.58, *p* value = 0.21, mean of co-culture *Neptunomonas* = 4.37 U/µL; *t* = -1.19, *p* value = 0.31). Stars indicate significant *p*-values (* = *p* < 0.05, ** = *p* < 0.01, *** = *p* < 0.001). (B) Compared to *V. natriegens* monocultures (left) the co-culture with *Alteromonas* (middle) leads to a decrease in *N*-acetyglucosaminidase activity of the substrate. The presence of *Neptunomonas* (right) has no effect. (Welch Two Sample t-test; N = 4; *N*-acetyglucosaminidase: mean of monoculture = 4.92 U/µL, mean of co-culture *Alteromonas* = 1.29 U/µL; *t* = 9.79, *p* value < 0.001, mean of co-culture *Neptunomonas* = 4.89 U/µL; *t* = 0.91, *p* value = 0.20. Chitobiosidases: mean of monoculture = 5.55 U/µL, mean of co-culture *Alteromonas* = 5.56 U/µL; *t* = -0.74, *p* value = 0.75, mean of co-culture *Neptunomonas* = 5.53 U/µL; *t* = 2.81, *p* value = 0.03. Endochitinases: mean of monoculture = 4.38 U/µL, mean of co-culture *Alteromonas* = 3.45 U/µL; *t* = 50.3, *p* value < 0.001, mean of co-culture *Neptunomonas* = 4.39 U/µL; *t* = -0.75, *p* value = 0.76). Stars indicate significant *p*-values (* = *p* < 0.05, ** = *p* < 0.01, *** = *p* < 0.001).

In summary, our findings show that secreted enzymes are themselves subject to degradation by microbial communities. This is further supported by the presence of proteases in the culture media of mono- and co-cultures of our focal strains (Figure S12 and S13). Therefore, chitin degradation activity by a community can be affected by non-chitin-degrading species in two ways: (i) through the modulation of chitinase gene expression in the degrader species, and (ii) through the degradation of extracellular chitinases for nutrient acquisition. These findings lead to new hypotheses about the role of species interactions on ecosystem processes carried out by microbial communities.

## Conclusion

Important questions in microbial ecology are (i) how are ecosystem functions determined by species composition? (ii) how do community level properties emerge from species interaction within microbial communities? and (iii) how do species from different trophic levels influence each other? We aimed to shed light on these questions using small synthetic communities of natural marine isolates that grow on the polymer chitin. Overall, we found striking effects of interspecies interactions on the behaviour of microbial species that perform key ecosystem functions, in our case chitin degradation. We found that species not directly involved in that function can still influence it in a highly specific manner. Concretely, we found that the presence of non-degrading species can influence chitin degradation activity by a multi-species community and that this happens through modulation of the expression of hydrolytic enzymes by the degrader species as well as through the degradation of these enzymes for use as nutrient source by community members. Hereby, our findings challenge the prevailing view that non-degraders do not contribute to overall biopolymer degradation activity at the community level (27) and provide mechanistic insights into these “downward” interactions.

Transcriptome profiling of degrader-cross-feeder co-cultures further revealed that the presence of the cross-feeder *Alteromonas* increased the expression of chitinases by the degrader *Vibrio* strains, which manifests on a community level as increases in chitin degradation activity. Furthermore, the presence of *Alteromonas* induced expression of general motility genes in *V. natriegens*, while simultaneously reducing expression of quorum sensing pathways. One possible explanation that connects these different findings is that *V. natriegens* responds with increased mobility to the presence of other species potentially as an avoidance strategy; as a consequence, more dispersed individual *V. natriegens* cells would need to excrete more enzymes in order to degrade and consume chitin. An alternative explanation is that the upregulation of chitinases leads to increased concentrations of chitobioses, which may then serve as chemoattractants in the environment. The chemoattractants may in turn induce genes related to motility such as flagellar assembly, which we observed. The causal connections and the mechanisms that underlie these correlations are still unknown. Our observations of gene expression changes in the presence of *Alteromonas* are in line with previous studies that show that growth behaviour on carbohydrate polymers such as aggregation or dispersal and the production of hydrolytic enzymes are tightly coupled (36). A study performed with various *Vibrio* species on the polymer alginate has shown that the presence of cross-feeders changes the aggregation of the degrading species (24). In combination, these findings suggest that secreted extracellular chitinases might provide two functions to the degraders. One is to generate nutrients via the cleavage of polymers, while the other is to generate a chemotactic gradient to fresh nutrient sources (21).

In addition to the modulation of gene expression in degrader species, non-degraders can also affect community-level chitin degradation more directly. We found that chitinases themselves can serve as a growth substrate. Furthermore, we found them to be degraded in an enzyme-specific manner and to different extents by different species, i.e., the depletion of specific chitinases in a community is dependent on community composition.

Our results highlight consequences of interspecies interactions on the behaviour of key microbial taxa that drive important ecosystem functions such as the degradation of complex polysaccharides. They reveal the importance of studying community level properties and cellular behaviour in the context of interspecies interactions.

## Material and Methods

### Bacterial strains, media and batch cultures

We used the wildtype strain *Vibrio natriegens* ATCC 14048, *Vibrio splendidus* sp. 1A01, *Vibrio splendidus* sp. 13B01, *Vibrio tasmaniensis* sp. 1F187, *Enterovibrio norvegicus* sp. 1F211, *Vibrio cyclitrophicus* sp. 1F273, *Vibrio rumoiensis* sp. 1S45, *Vibrio lentus* sp. 5F79, *Vibrio kanaloae* sp. 5S149, *Vibrio ordalii* sp. FS144, *Aliivibrio fischeri*, sp. ZF211 as our degrader species (25, 27, 37). *Alteromonas macleodii* sp. 4B03 *Alteromonas macleodii* sp. A, *Neptunomonas phycophila* sp. 3B05, *Shewanella* sp. A2R10, *Colwellia psychrerythraea* sp. C2M11, *Marinobacter* sp. D2M19 as our non-degrader species (27).

Strains were inoculated from cryo-culture at -80 °C into 3 mL Marine Broth (MB, Difco 2216) and grown over night at 25 °C and 200 rpm. 1mL of these cell cultures was centrifuged (13000 rpm for 2 min) in a 1.5ml microfuge tube. Supernatant was discarded and the cells were washed with 1ml of MBL minimal medium (38) without carbon source in order to remove excess carbon. Cells were centrifuged again and the cell pellet was resuspended in 1mL of carbon depleted MBL adjusted to an OD600 of 0.002.

### Growth assays

From the above described cultures, cells were used for experiments in MBL minimal medium with 0.1% (weight/volume) Pentaacetyl-Chitopentaose (Megazyme, Ireland). The carbon source was added to the MBL minimum medium and filter sterilized using 0.22μm Surfactant-Free Cellulose Acetate filters (Corning, USA). A total of 10uL of cell culture was added to 190uL of MBL containing 0.1% Chitopentaose (w/v). This yielded a final starting OD of 0.0001. Cultures were grown in 96-well plates for 48h in a plate reader (Eon, BioTek) at 25°C.

### Chitinase assays

Cell free supernatants were generated by sterile filtering cultures using a multi-well filter plate (AcroPrep) into a fresh 96-well plate. Chitinase activity of cell free supernatants was measured using a commercially available fluorometric chitinase assay kit (Sigma-Aldrich) following the protocol. In short, (i) β-N-acetylglucosaminidase, (ii) chitobiosidase, and (iii) endochitinase activity was measured using 4-methylumbelliferyl N-acetyl-β-d-glucosaminide, 4-methylumbelliferyl N,N′-diacetyl-β-d-chitobioside, and 4-methylumbelliferyl β-d-N,N′,N″-triacetylchitotriose substrates, respectively. 10 μl of cell free supernatant was added to the 90 μl each of the three substrate solutions and incubated in the dark for 40min at 25 °C. Thereafter, the reactions were halted by adding 200 μl of Stop Solution (39). Fluorescence of released 4-methylumbelliferone (4MU) was measured (Excitation 360 nm, Emission 450 nm) in a plate reader (Synergy MX, Biotek). One unit of chitinase activity releases 1 μmol of 4MU from the appropriate substrate per minute. The chitinase activity was calculated using a single standard concentration (1.9 nmol/mL) and the following equation:

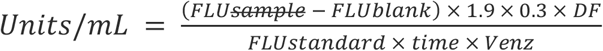

Where FLU – fluorescence of the sample, FLUblank – fluorescence of the blank, 0.3 – final reaction volume in mL, DF – enzyme dilution factor, FLUstandard – fluorescence of the Standard Solution, Time – minutes, Venz – volume of the sample in mL.

### Chitinase substrate assay

From the above described cultures, cells were used for experiments in MBL minimal medium with 0.04% (weight/volume) chitinase from Trichoderma viride (Sigma-Aldrich, C6202). The lyophilized enzyme was added to the MBL minimum medium and filter sterilized using 0.22μm Surfactant-Free Cellulose Acetate filters (Corning, USA). A total of 10uL of cell culture was added to 190uL of MBL containing 0.04% chitinase (w/v). This yielded a final starting OD of 0.0001. Cultures were grown in 96-well plates for 48h in a plate reader (Eon, BioTek) at 25°C.

### Protease Assay

Cell free supernatants of cultures grown on chitinase as a sole carbon source were generated by sterile filtering cultures using a multi-well filter plate (AcroPrep) into a fresh 96-well plate. Protease activity of cell free supernatants was measured using the commercially available Pierce Colorimetric Protease Assay kit (Thermo FisherScientific; 23263). Lyophilized succinylated casein was resuspended in MBL to a concentration of 2 mg/mL. 50 μL of supernatant was added to 100 μL of succinylated casein and incubated at 25 °C for 1 h in a 96-well plate. 50 μL of 0.033% TNBSA was added, and samples were incubated for additional 20 min at room temperature. Absorbance (450 nm) was measured in a 96-well plate and corrected by subtracting a control containing a buffer instead of supernatant. The final protease activity was calculated by subtracting the corrected succinylated casein without supernatant signal from the supernatant signal (40).

### Statistical analysis

Growth curves were analyzed in Python (v3.8) using the AMiGA software (41). All statistical analysis was performed R Studio v2021.09.1 (42) with R v4.1.2. Each boxplot figure depicts median of the corresponding value. Measures of effect size are represented by the R^2^. Statistically significant differences in chitinase activity between mono- and co-cultures were calculated using ANOVA models with the LmerTest package v 0.99.45 (43) with the equation (Chitinase activity/yield) ∼ Crossfeeder + Replicate. The Fisher-LSD Posthoc test was performed using the same software. For the number of replicates please refer to the supplementary materials. The heatmaps were generated using the ggplot2 package v3.3.3.9000 (44). In order to generate the color gradients, values were scaled using the mutate function in the dplyr packages v.1.0.5 (45).

### RNA extraction

For RNA extraction, cells were grown as described above and harvested at exponential growth phase. 4 ml of cell culture was centrifuged at 5000xg for 10 min. Supernatant was removed and the cell pellets were resuspended in 1 ml of RNAprotect Bacteria Reagent (Qiagen) and stored -20 until RNA extraction. To extract the total RNA, samples were thawed and the suspension was centrifuged at 5000xg for 10 min, the supernatant was carefully removed. To extract the total RNA of the frozen cell suspensions, we followed the protocol of the RNeasy Protect Bacteria Kit (Qiagen) combined with on-column DNase digestion using the RNase-Free DNase Set (Qiagen). Cells were incubated 15 min at room temperature with 1 mg/ml lysozyme (Sigma) for lysis. The isolated total RNA was stored at -80 °C until sending it for sequencing.

### RNA sequencing

RNA sequencing was performed by the Oxford Genomics Center (OGC, Oxford, UK). The sequencing package included rRNA depletion. All but one sample met the quality requirements of OGC (>100 ng/ul RNA concentration). The sample that did not meet the requirements was a monoculture of *V. natriegens, therefore* the corresponding replicates of the co-cultures of *Alteromonas* and *Neptunomonas* were disregarded from the downstream analysis. Ribodepletions using NEB bacterial probes before conversion to cDNA was performed. Second strand cDNA synthesis did incorporate dUTP. The cDNA was end-repaired, A-tailed and adapter-ligated. Prior to amplification, samples did undergo uridine digestion. The prepared libraries were size selected, multiplexed and QC’ed before paired end sequencing over one unit of a flow cell. Sequencing was conducted on a NovaSeq6000 system (Illumina).

### RNA-seq: Pre-processing

Preprocessing of the raw reads was carried out as follows: A quality control was performed with FastQC v0.11.9 (46) and Trimmomatic v0.38 (47); the high-quality reads were mapped to the reference genome (RefSeq assembly accession GCF_000006905.1) with Bowtie2 v2.3.5.1 (48); binarisation, sorting, and indexing were done with Samtools v1.10 (49); gene counts were computed with the featureCount function of Subread v2.0.1 (50).

### RNA-seq: differential expression analysis

Differential expression was calculated using DESeq2 v1.34.0 (51) in R Studio v2021.09.1 (42) with R v4.1.2. For the analysis, all rRNA and tRNA genes were excluded. Genes were considered to be differentially expressed if they had an FDR < 0.05 and a log2-Fold change (log2-FC) either higher than 0.6 or lower than -0.6. Heatmaps of genes with most significant expression changes were computed by first transforming the raw counts with the variance stabilizing transformation (VST) function of DESeq2 and then using the pheatmap function of the Pheatmap package v1.0.12 (52).

### RNA-seq: functional analysis

For annotations of KEGG orthology (53) BlastKOALA v2.2 (54) was used to assign annotations. eggNOG v5.0 (55) was used to assign the Cluster of Orthologous Groups (COG) annotations (56) for all genes. Dbcan2 (57) was used to annotate carbohydrate active enzymes according to the Carbohydrate-Active Enzymes (CAZy) Database (58).

## Acknowledgments

We thank Roman Stocker and Shini Sunagawa for advice for data analysis and for fruitful scientific discussions. We also thank the members of the Simons Foundation Collaboration on Principles of Microbial Ecosystems collaboration for feedback on the project.

