## Supplementary Figures for "Effects of interspecies interactions on marine community ecosystem function"

### Supplementary Material

#### Annotation of chitinases

Microorganisms often contain chitinase genes that encode for functionally similar enzymes (Wang et al., 2019). The biochemical classification of these enzymes is carried out in two ways; with their annotation to an Enzyme Commission number (EC number) based on the enzyme's functionality (Enzyme nomenclature, 1992) and by grouping the enzymes into glycoside hydrolase (GH) family based on the enzymes amino acid sequence similarities and fold structure (Henrissat et al., 1995).

The classification into the EC number is based on the chemical reaction that an enzyme catalyses. In cases where different enzymes i.e. from different organisms carry out the same reaction they are given the same EC number. It is an important classification when enzymes with the same function have different evolutionary origins and therefore different protein folds (Hunt et al., 2008). The classification into GH families is based on structural similarities. This approach is important because in genetically close organisms enzymatic properties can be structurally highly conserved but functionally diverse (Hehemann et al., 2016).

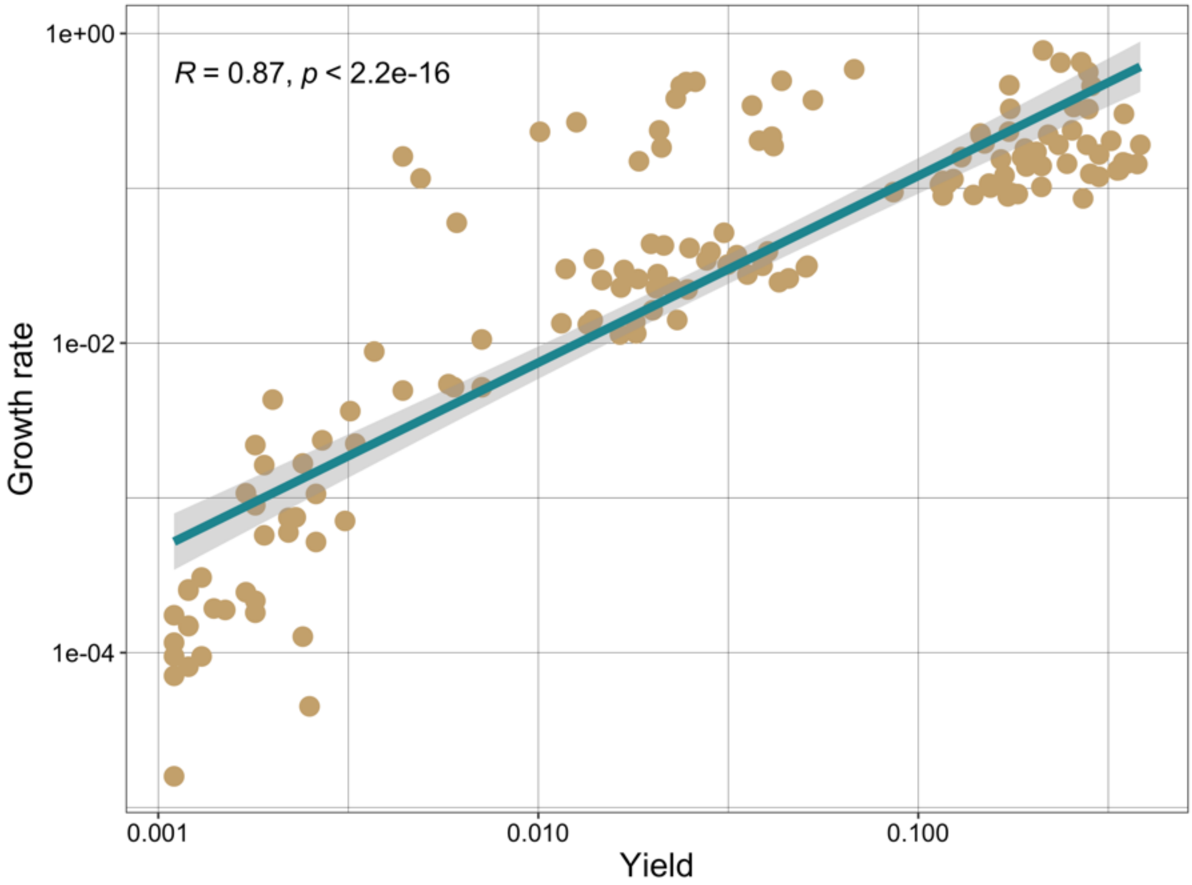


**Figure S1: Yield and growth rate of degrader monocultures are coupled.**

Growth rate is positively correlated with yield of degrader monocultures. Growth rate of individual degrader monocultures as a function of their yield (dots).

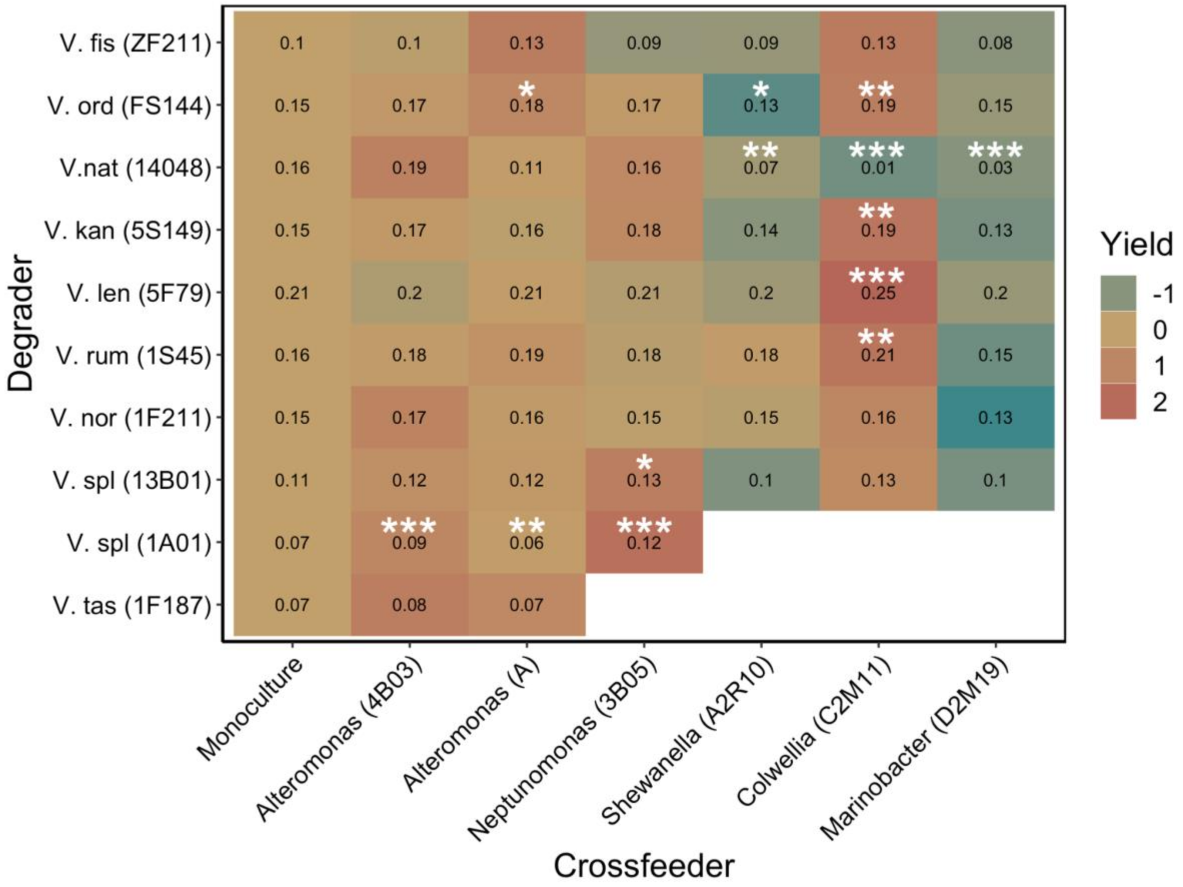


**Figure S2: Cross-feeders influence community level yield**

Heatmap of yield for various degraders (y-axis) in monoculture (first column) or in co-culture with different cross-feeders (x-axis). Numbers indicate total yield of the cultures. Colours represent normalised differences in yield in co-culture compared to respective degrader monoculture. Yellow colours indicate little to no change in yield. Green colours indicate a reduced yield. Red colours represent an increase in yield. Stars represent significant changes in yield between the monoculture and respective co-culture. * = $p < 0.05$, ** = $p < 0.01$, *** = $p < 0.001$. A detailed summary of the ANOVA results can be found in the supplementary materials (Table S1).

| Degrader \ Crossfeeder | Alteromonas (4B03) | Alteromonas (A) | Neptunomonas (3B05) | Shewanella (A2R10) | Colwellia (C2M11) | Marinobacter (D2M19) |
|---|---|---|---|---|---|---|
| V. fis (ZF211) | d = 0, p = 0.953, cl = -0.05, cu = 0.04 | d = 0.03, p = 0.243, cl = -0.02, cu = 0.07 | d = -0.01, p = 0.548, cl = -0.06, cu = 0.03 | d = -0.01, p = 0.558, cl = -0.06, cu = 0.03 | d = 0.03, p = 0.256, cl = -0.02, cu = 0.08 | d = -0.02, p = 0.444, cl = -0.06, cu = 0.03 |
| V. ord (FS144) | d = 0.01, p = 0.252, cl = -0.01, cu = 0.04 | d = 0.03, p = 0.026, cl = 0, cu = 0.05 | d = 0.01, p = 0.291, cl = -0.01, cu = 0.03 | d = -0.03, p = 0.02, cl = -0.05, cu = 0 | d = 0.03, p = 0.008, cl = 0.01, cu = 0.06 | d = -0.01, p = 0.446, cl = -0.03, cu = 0.01 |
| V.nat (14048) | d = 0.02, p = 0.346, cl = -0.03, cu = 0.07 | d = -0.05, p = 0.086, cl = -0.1, cu = 0.01 | d = 0, p = 0.924, cl = -0.05, cu = 0.05 | d = -0.1, p = 0.001, cl = -0.15, cu = -0.04 | d = -0.15, p = 0, cl = -0.21, cu = -0.09 | d = -0.13, p = 0, cl = -0.19, cu = -0.07 |
| V. kan (5S149) | d = 0.01, p = 0.324, cl = -0.01, cu = 0.04 | d = 0, p = 0.735, cl = -0.02, cu = 0.03 | d = 0.02, p = 0.053, cl = 0, cu = 0.05 | d = -0.01, p = 0.25, cl = -0.04, cu = 0.01 | d = 0.04, p = 0.005, cl = 0.01, cu = 0.07 | d = -0.02, p = 0.12, cl = -0.04, cu = 0.01 |
| V. len (5F79) | d = 0, p = 0.55, cl = -0.02, cu = 0.01 | d = 0, p = 0.416, cl = -0.01, cu = 0.02 | d = 0, p = 0.732, cl = -0.01, cu = 0.01 | d = -0.01, p = 0.186, cl = -0.02, cu = 0 | d = 0.04, p = 0, cl = 0.03, cu = 0.05 | d = -0.01, p = 0.113, cl = -0.02, cu = 0 |
| V. rum (1S45) | d = 0.02, p = 0.15, cl = -0.01, cu = 0.05 | d = 0.03, p = 0.061, cl = 0, cu = 0.06 | d = 0.01, p = 0.317, cl = -0.01, cu = 0.04 | d = 0.02, p = 0.142, cl = -0.01, cu = 0.05 | d = 0.05, p = 0.004, cl = 0.02, cu = 0.08 | d = -0.01, p = 0.603, cl = -0.04, cu = 0.02 |
| V. nor (1F211) | d = 0.01, p = 0.387, cl = -0.02, cu = 0.04 | d = 0, p = 0.862, cl = -0.02, cu = 0.03 | d = 0, p = 0.904, cl = -0.03, cu = 0.03 | d = 0, p = 0.844, cl = -0.03, cu = 0.02 | d = 0.01, p = 0.49, cl = -0.02, cu = 0.04 | d = -0.02, p = 0.1, cl = -0.05, cu = 0 |
| V. spl (13B01) | d = 0.02, p = 0.148, cl = -0.01, cu = 0.04 | d = 0.01, p = 0.268, cl = -0.01, cu = 0.04 | d = 0.03, p = 0.026, cl = 0, cu = 0.05 | d = -0.01, p = 0.496, cl = -0.03, cu = 0.02 | d = 0.02, p = 0.115, cl = -0.01, cu = 0.05 | d = -0.01, p = 0.421, cl = -0.03, cu = 0.01 |
| V. spl (1A01) | d = 0.02, p = 0, cl = 0.01, cu = 0.02 | d = -0.01, p = 0.002, cl = -0.02, cu = 0 | d = 0.04, p = 0, cl = 0.04, cu = 0.05 | | | |
| V. tas (1F187) | d = 0, p = 0.58, cl = -0.01, cu = 0.02 | d = 0, p = 0.822, cl = -0.02, cu = 0.01 | | | | |

**Table S1: ANOVA results yield (related to Figure S2)**

ANOVA table of yield differences for combinations of various degraders (y-axis) in co-culture with different cross-feeders (x-axis). Posthoc test for multiple comparisons of means using Fisher LSD of a two-way ANOVA (with the factors culture-type and day of the experiment) revealed these scores. Shown are calculated differences in yield between mono- and co-culture (d), the associated *p* – value (p), and the lower and upper 95% confidence intervals (cl, cu). In cases where the rounded digits indicate p = 0, p < 0.0005. In cases where the rounded digits indicate d = 0, d < 0.005.

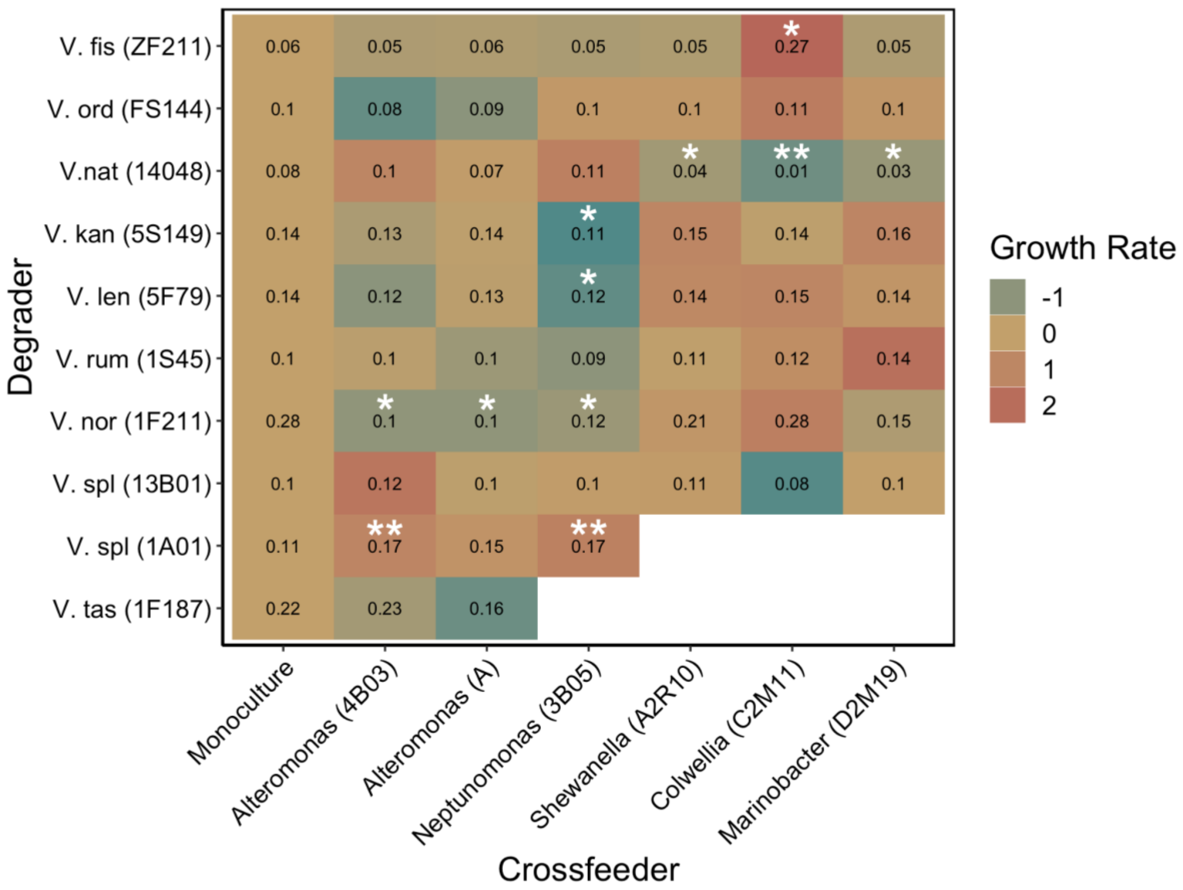


**Figure S3: Cross-feeders influence community level growth rate**

Heatmap of growth rate for various degraders (y-axis) in monoculture (first column) or in co-culture with different cross-feeders (x-axis). Numbers indicate maximal growth rate of the cultures. Colours represent normalised differences in growth rate in co-culture compared to respective degrader monoculture. Yellow colours indicate little to no change in growth rate. Green colours indicate a reduced growth rate. Red colours represent an increase in growth rate. Stars represent significant changes in growth rate between the monoculture and respective co-culture. * = $p < 0.05$, ** = $p < 0.01$, *** = $p < 0.001$. A detailed summary of the ANOVA results can be found in the supplementary materials (Table S2).

| Degrader \ Crossfeeder | Alteromonas (4B03) | Alteromonas (A) | Neptunomonas (3B05) | Shewanella (A2R10) | Colwellia (C2M11) | Marinobacter (D2M19) |
|---|---|---|---|---|---|---|
| V. fis (ZF211) | d = -0.01 p = 0.881<br>cl = -0.17 cu = 0.15 | d = -0.01 p = 0.918<br>cl = -0.17 cu = 0.15 | d = -0.02 p = 0.845<br>cl = -0.18 cu = 0.14 | d = -0.01 p = 0.906<br>cl = -0.17 cu = 0.15 | d = 0.2 p = 0.024<br>cl = 0.03 cu = 0.38 | d = -0.01 p = 0.884<br>cl = -0.17 cu = 0.15 |
| V. ord (FS144) | d = -0.01 p = 0.323<br>cl = -0.04 cu = 0.01 | d = -0.01 p = 0.513<br>cl = -0.04 cu = 0.02 | d = 0 p = 0.907<br>cl = -0.03 cu = 0.03 | d = 0 p = 0.901<br>cl = -0.03 cu = 0.03 | d = 0.01 p = 0.487<br>cl = -0.02 cu = 0.04 | d = 0 p = 0.869<br>cl = -0.03 cu = 0.03 |
| V.nat (14048) | d = 0.02 p = 0.172<br>cl = -0.01 cu = 0.06 | d = -0.01 p = 0.597<br>cl = -0.05 cu = 0.03 | d = 0.03 p = 0.053<br>cl = 0 cu = 0.06 | d = -0.04 p = 0.043<br>cl = -0.07 cu = 0 | d = -0.07 p = 0.001<br>cl = -0.11 cu = -0.03 | d = -0.04 p = 0.019<br>cl = -0.08 cu = -0.01 |
| V. kan (5S149) | d = -0.01 p = 0.345<br>cl = -0.04 cu = 0.02 | d = -0.01 p = 0.579<br>cl = -0.04 cu = 0.02 | d = -0.04 p = 0.015<br>cl = -0.07 cu = -0.01 | d = 0.01 p = 0.471<br>cl = -0.02 cu = 0.04 | d = -0.01 p = 0.606<br>cl = -0.04 cu = 0.02 | d = 0.01 p = 0.449<br>cl = -0.02 cu = 0.04 |
| V. len (5F79) | d = -0.01 p = 0.101<br>cl = -0.03 cu = 0 | d = -0.01 p = 0.513<br>cl = -0.02 cu = 0.01 | d = -0.02 p = 0.02<br>cl = -0.04 cu = 0 | d = 0.01 p = 0.563<br>cl = -0.01 cu = 0.02 | d = 0.01 p = 0.514<br>cl = -0.01 cu = 0.03 | d = 0 p = 0.97<br>cl = -0.02 cu = 0.02 |
| V. rum (1S45) | d = 0.01 p = 0.772<br>cl = -0.05 cu = 0.07 | d = 0 p = 0.981<br>cl = -0.06 cu = 0.06 | d = -0.01 p = 0.857<br>cl = -0.07 cu = 0.05 | d = 0.01 p = 0.73<br>cl = -0.05 cu = 0.07 | d = 0.02 p = 0.47<br>cl = -0.04 cu = 0.09 | d = 0.04 p = 0.156<br>cl = -0.02 cu = 0.1 |
| V. nor (1F211) | d = -0.17 p = 0.022<br>cl = -0.32 cu = -0.03 | d = -0.17 p = 0.022<br>cl = -0.32 cu = -0.03 | d = -0.16 p = 0.035<br>cl = -0.3 cu = -0.01 | d = -0.07 p = 0.322<br>cl = -0.22 cu = 0.07 | d = 0 p = 0.991<br>cl = -0.16 cu = 0.16 | d = -0.13 p = 0.078<br>cl = -0.28 cu = 0.02 |
| V. spl (13B01) | d = 0.02 p = 0.277<br>cl = -0.02 cu = 0.06 | d = 0 p = 0.999<br>cl = -0.04 cu = 0.04 | d = 0 p = 0.865<br>cl = -0.04 cu = 0.04 | d = 0 p = 0.849<br>cl = -0.03 cu = 0.04 | d = -0.02 p = 0.337<br>cl = -0.06 cu = 0.02 | d = 0 p = 0.93<br>cl = -0.04 cu = 0.04 |
| V. spl (1A01) | d = 0.06 p = 0.007<br>cl = 0.02 cu = 0.1 | d = 0.04 p = 0.076<br>cl = 0 cu = 0.09 | d = 0.06 p = 0.009<br>cl = 0.02 cu = 0.11 | | | |
| V. tas (1F187) | d = 0.01 p = 0.85<br>cl = -0.11 cu = 0.14 | d = -0.06 p = 0.314<br>cl = -0.18 cu = 0.06 | | | | |

**Table S2: ANOVA results growth rate (related to Figure S3)**

ANOVA table of differences in growth rate for combinations of various degraders (y-axis) in co-culture with different cross-feeders (x-axis). Posthoc test for multiple comparisons of means using Fisher LSD of a two-way ANOVA (with the factors culture-type and day of the experiment) revealed these scores. Shown are calculated differences in growth rate between mono- and co-culture (d), the associated *p* – value (p), and the lower and upper 95% confidence intervals (cl, cu). In cases where the rounded digits indicate p = 0, p < 0.0005. In cases where the rounded digits indicate d = 0, d < 0.005.

| Degrader \ Crossfeeder | Alteromonas (4B03) | Alteromonas (A) | Neptunomonas (3B05) | Shewanella (A2R10) | Colwellia (C2M11) | Marinobacter (D2M19) |
|---|---|---|---|---|---|---|
| V. fis (ZF211) | d = 0.33; p = 0.013; cl = 0.07; cu = 0.58 | d = 0.2; p = 0.117; cl = -0.05; cu = 0.46 | d = 0.05; p = 0.719; cl = -0.21; cu = 0.3 | d = -0.2; p = 0.117; cl = -0.46; cu = 0.05 | d = 0.09; p = 0.516; cl = -0.19; cu = 0.36 | d = -0.12; p = 0.359; cl = -0.37; cu = 0.14 |
| V. ord (FS144) | d = 0.21; p = 0.065; cl = -0.01; cu = 0.43 | d = -0.06; p = 0.59; cl = -0.28; cu = 0.16 | d = -0.28; p = 0.017; cl = -0.5; cu = -0.05 | d = -0.09; p = 0.419; cl = -0.31; cu = 0.13 | d = -0.03; p = 0.805; cl = -0.27; cu = 0.21 | d = 0.01; p = 0.941; cl = -0.21; cu = 0.23 |
| V.nat (14048) | d = 0.95; p = 0; cl = 0.53; cu = 1.37 | d = 1.09; p = 0; cl = 0.62; cu = 1.57 | d = 0.08; p = 0.697; cl = -0.34; cu = 0.5 | d = -0.13; p = 0.594; cl = -0.6; cu = 0.35 | d = -0.22; p = 0.403; cl = -0.73; cu = 0.3 | d = -0.09; p = 0.714; cl = -0.56; cu = 0.39 |
| V. kan (5S149) | d = -0.29; p = 0.308; cl = -0.86; cu = 0.28 | d = -0.7; p = 0.018; cl = -1.27; cu = -0.13 | d = -0.72; p = 0.014; cl = -1.29; cu = -0.15 | d = -0.33; p = 0.24; cl = -0.9; cu = 0.23 | d = -0.52; p = 0.096; cl = -1.15; cu = 0.1 | d = -0.06; p = 0.838; cl = -0.63; cu = 0.51 |
| V. len (5F79) | d = -0.6; p = 0.412; cl = -2.07; cu = 0.87 | d = -1.09; p = 0.141; cl = -2.56; cu = 0.38 | d = -0.29; p = 0.688; cl = -1.76; cu = 1.18 | d = -0.47; p = 0.521; cl = -1.94; cu = 1 | d = 0.01; p = 0.987; cl = -1.6; cu = 1.62 | d = -0.25; p = 0.729; cl = -1.72; cu = 1.22 |
| V. rum (1S45) | d = -1.15; p = 0.194; cl = -2.92; cu = 0.61 | d = -1.32; p = 0.137; cl = -3.09; cu = 0.44 | d = -0.65; p = 0.463; cl = -2.41; cu = 1.12 | d = -1.75; p = 0.052; cl = -3.52; cu = 0.01 | d = -1.15; p = 0.229; cl = -3.06; cu = 0.75 | d = -1.32; p = 0.138; cl = -3.09; cu = 0.44 |
| V. nor (1F211) | d = -0.62; p = 0.106; cl = -1.38; cu = 0.14 | d = -0.54; p = 0.156; cl = -1.3; cu = 0.22 | d = -0.94; p = 0.017; cl = -1.7; cu = -0.18 | d = -0.22; p = 0.565; cl = -0.97; cu = 0.54 | d = -0.17; p = 0.687; cl = -1.02; cu = 0.68 | d = -0.02; p = 0.95; cl = -0.78; cu = 0.73 |
| V. spl (13B01) | d = 0.48; p = 0.222; cl = -0.3; cu = 1.27 | d = 0.36; p = 0.365; cl = -0.43; cu = 1.15 | d = -0.99; p = 0.015; cl = -1.78; cu = -0.2 | d = -0.64; p = 0.107; cl = -1.43; cu = 0.14 | d = -0.62; p = 0.151; cl = -1.47; cu = 0.23 | d = -0.35; p = 0.371; cl = -1.14; cu = 0.44 |
| V. spl (1A01) | d = 1.62; p = 0.006; cl = 0.5; cu = 2.73 | d = 2.95; p = 0; cl = 1.66; cu = 4.23 | d = -0.76; p = 0.239; cl = -2.04; cu = 0.53 | | | |
| V. tas (1F187) | d = 1.16; p = 0.066; cl = -0.08; cu = 2.41 | d = 0.98; p = 0.115; cl = -0.26; cu = 2.23 | | | | |

**Table S3: ANOVA results chitobiosidase (related to Figure 3)**

ANOVA table of differences in chitobiosidase activity for combinations of various degraders (y-axis) in co-culture with different cross-feeders (x-axis). Posthoc test for multiple comparisons of means using Fisher LSD of a two-way ANOVA (with the factors culture-type and day of the experiment) revealed these scores. Shown are calculated differences in chitobiosidase activity mono- and co-culture (d), the associated *p* – value (p), and the lower and upper 95% confidence intervals (cl, cu). In cases where the rounded digits indicate p = 0, p < 0.0005. In cases where the rounded digits indicate d = 0, d < 0.005.

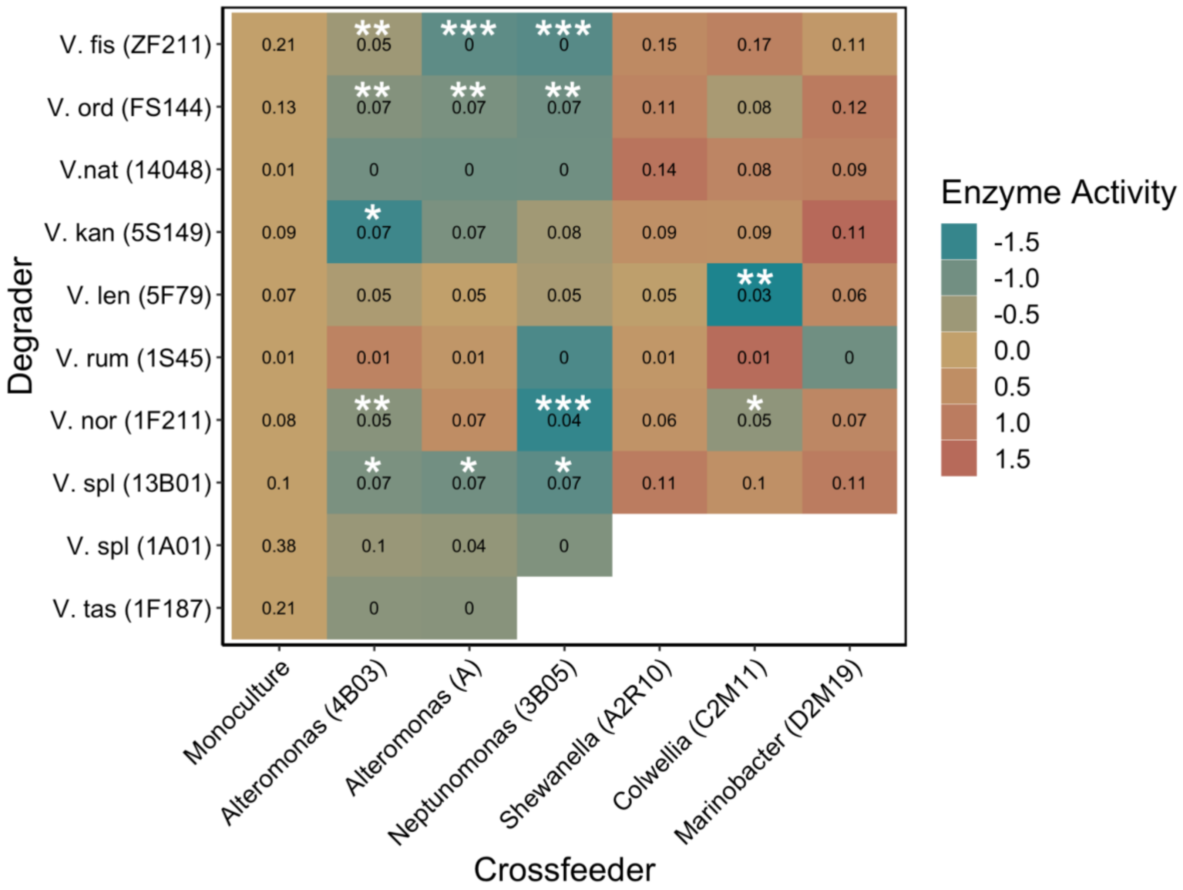


**Figure S4: Cross-feeders reduce community level activity of N-acetyglucosaminidases**

Heatmap of N-acetyglucosaminidase activity for various degraders (y-axis) in monoculture (first column) or in co-culture with different cross-feeders (x-axis). Numbers indicate enzyme activity of the cultures. Colours represent normalised differences in enzyme activity in co-culture compared to respective degrader monoculture. Yellowish colours indicate little to no change in enzyme activity. Greenish colours indicate a reduced enzyme activity. Reddish colours represent an increase in enzyme activity. Stars represent significant changes in growth between the monoculture and respective co-culture. * = $p < 0.05$, ** = $p < 0.01$, *** = $p < 0.001$. A detailed summary of the ANOVA results can be found in the supplementary materials (Table S4).

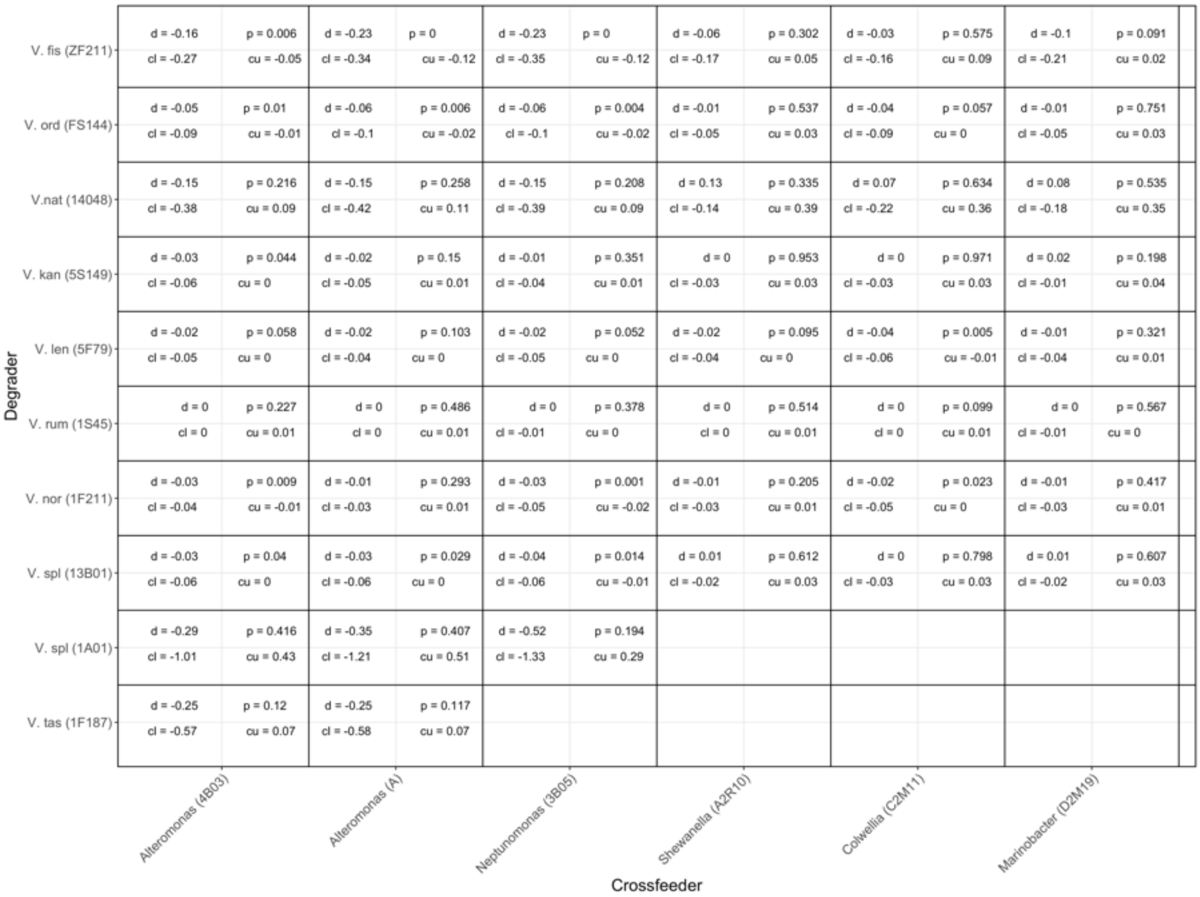

| Degrader \ Crossfeeder | Alteromonas (4B03) | Alteromonas (A) | Neptunomonas (3B05) | Shewanella (A2R10) | Colwellia (C2M11) | Marinobacter (D2M19) |
|---|---|---|---|---|---|---|
| V. fis (ZF211) | d = -0.16 p = 0.006<br>cl = -0.27 cu = -0.05 | d = -0.23 p = 0<br>cl = -0.34 cu = -0.12 | d = -0.23 p = 0<br>cl = -0.35 cu = -0.12 | d = -0.06 p = 0.302<br>cl = -0.17 cu = 0.05 | d = -0.03 p = 0.575<br>cl = -0.16 cu = 0.09 | d = -0.1 p = 0.091<br>cl = -0.21 cu = 0.02 |
| V. ord (FS144) | d = -0.05 p = 0.01<br>cl = -0.09 cu = -0.01 | d = -0.06 p = 0.006<br>cl = -0.1 cu = -0.02 | d = -0.06 p = 0.004<br>cl = -0.1 cu = -0.02 | d = -0.01 p = 0.537<br>cl = -0.05 cu = 0.03 | d = -0.04 p = 0.057<br>cl = -0.09 cu = 0 | d = -0.01 p = 0.751<br>cl = -0.05 cu = 0.03 |
| V.nat (14048) | d = -0.15 p = 0.216<br>cl = -0.38 cu = 0.09 | d = -0.15 p = 0.258<br>cl = -0.42 cu = 0.11 | d = -0.15 p = 0.208<br>cl = -0.39 cu = 0.09 | d = 0.13 p = 0.335<br>cl = -0.14 cu = 0.39 | d = 0.07 p = 0.634<br>cl = -0.22 cu = 0.36 | d = 0.08 p = 0.535<br>cl = -0.18 cu = 0.35 |
| V. kan (5S149) | d = -0.03 p = 0.044<br>cl = -0.06 cu = 0 | d = -0.02 p = 0.15<br>cl = -0.05 cu = 0.01 | d = -0.01 p = 0.351<br>cl = -0.04 cu = 0.01 | d = 0 p = 0.953<br>cl = -0.03 cu = 0.03 | d = 0 p = 0.971<br>cl = -0.03 cu = 0.03 | d = 0.02 p = 0.198<br>cl = -0.01 cu = 0.04 |
| V. len (5F79) | d = -0.02 p = 0.058<br>cl = -0.05 cu = 0 | d = -0.02 p = 0.103<br>cl = -0.04 cu = 0 | d = -0.02 p = 0.052<br>cl = -0.05 cu = 0 | d = -0.02 p = 0.095<br>cl = -0.04 cu = 0 | d = -0.04 p = 0.005<br>cl = -0.06 cu = -0.01 | d = -0.01 p = 0.321<br>cl = -0.04 cu = 0.01 |
| V. rum (1S45) | d = 0 p = 0.227<br>cl = 0 cu = 0.01 | d = 0 p = 0.486<br>cl = 0 cu = 0.01 | d = 0 p = 0.378<br>cl = -0.01 cu = 0 | d = 0 p = 0.514<br>cl = 0 cu = 0.01 | d = 0 p = 0.099<br>cl = 0 cu = 0.01 | d = 0 p = 0.567<br>cl = -0.01 cu = 0 |
| V. nor (1F211) | d = -0.03 p = 0.009<br>cl = -0.04 cu = -0.01 | d = -0.01 p = 0.293<br>cl = -0.03 cu = 0.01 | d = -0.03 p = 0.001<br>cl = -0.05 cu = -0.02 | d = -0.01 p = 0.205<br>cl = -0.03 cu = 0.01 | d = -0.02 p = 0.023<br>cl = -0.05 cu = 0 | d = -0.01 p = 0.417<br>cl = -0.03 cu = 0.01 |
| V. spl (13B01) | d = -0.03 p = 0.04<br>cl = -0.06 cu = 0 | d = -0.03 p = 0.029<br>cl = -0.06 cu = 0 | d = -0.04 p = 0.014<br>cl = -0.06 cu = -0.01 | d = 0.01 p = 0.612<br>cl = -0.02 cu = 0.03 | d = 0 p = 0.798<br>cl = -0.03 cu = 0.03 | d = 0.01 p = 0.607<br>cl = -0.02 cu = 0.03 |
| V. spl (1A01) | d = -0.29 p = 0.416<br>cl = -1.01 cu = 0.43 | d = -0.35 p = 0.407<br>cl = -1.21 cu = 0.51 | d = -0.52 p = 0.194<br>cl = -1.33 cu = 0.29 | | | |
| V. tas (1F187) | d = -0.25 p = 0.12<br>cl = -0.57 cu = 0.07 | d = -0.25 p = 0.117<br>cl = -0.58 cu = 0.07 | | | | |

**Table S4: ANOVA results N-acetyglucosaminidase (Figure S4)**

ANOVA table of differences in N-acetyglucosaminidase activity for combinations of various degraders (y-axis) in co-culture with different cross-feeders (x-axis). Posthoc test for multiple comparisons of means using Fisher LSD of a two-way ANOVA (with the factors culture-type and day of the experiment) revealed these scores. Shown are calculated differences in N-acetyglucosaminidase activity mono- and co-culture (d), the associated *p*-value (p), and the lower and upper 95% confidence intervals (cl, cu). In cases where the rounded digits indicate p = 0, p < 0.0005. In cases where the rounded digits indicate d = 0, d < 0.005.

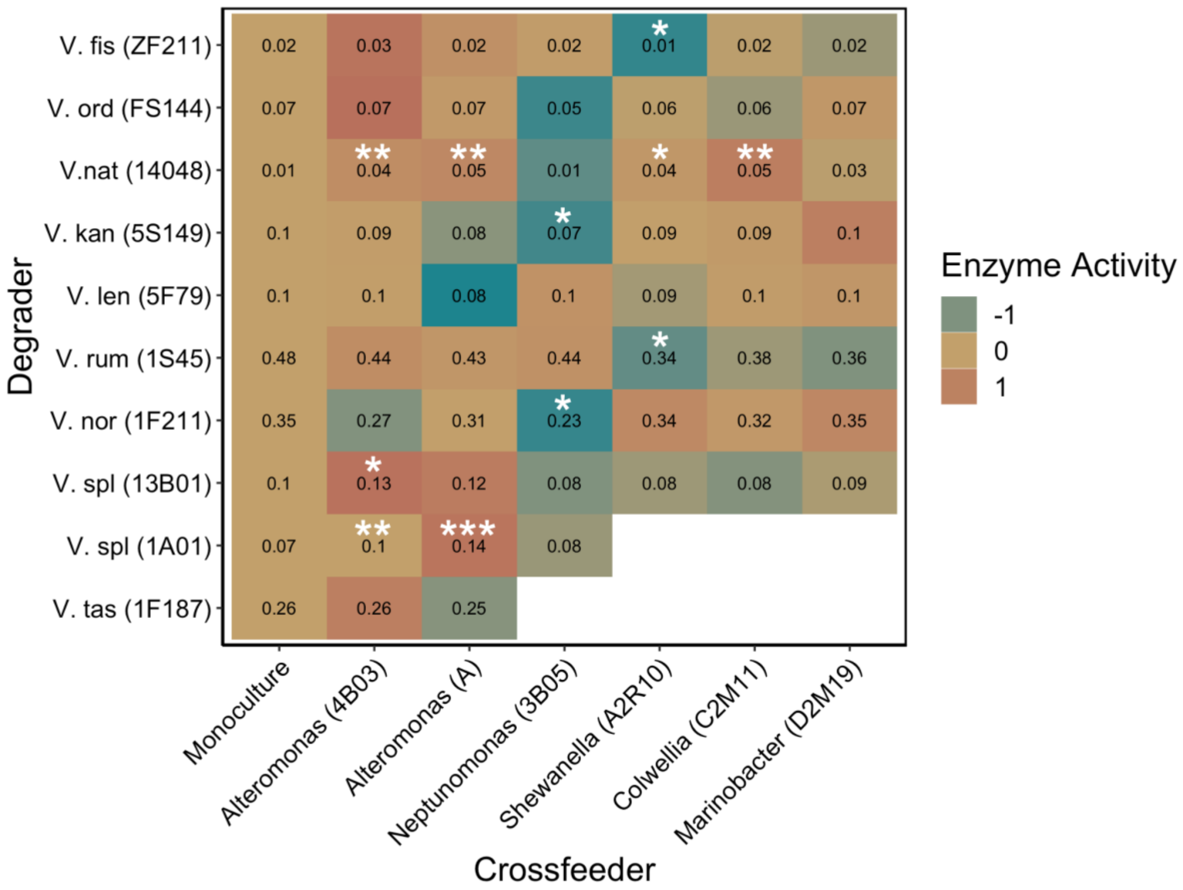


**Figure S5: Cross-feeders influence community level activity of endochitinases**

Heatmap of endochitinase activity for various degraders (y-axis) in monoculture (first column) or in co-culture with different cross-feeders (x-axis). Numbers indicate enzyme activity of the cultures. Colours represent normalised differences in enzyme activity in co-culture compared to respective degrader monoculture. Yellowish colours indicate little to no change in enzyme activity. Greenish colours indicate a reduced enzyme activity. Reddish colours represent an increase in enzyme activity. Stars represent significant changes in growth between the monoculture and respective co-culture. * = $p < 0.05$, ** = $p < 0.01$, *** = $p < 0.001$. A detailed summary of the ANOVA results can be found in Table S5.

| Degrader \ Crossfeeder | Alteromonas (4B03) | Alteromonas (A) | Neptunomonas (3B05) | Shewanella (A2R10) | Colwellia (C2M11) | Marinobacter (D2M19) |
|---|---|---|---|---|---|---|
| V. fis (ZF211) | d = 0.01; p = 0.251; cl = 0; cu = 0.01 | d = 0; p = 0.899; cl = -0.01; cu = 0.01 | d = 0; p = 0.759; cl = -0.01; cu = 0.01 | d = -0.01; p = 0.019; cl = -0.02; cu = 0 | d = 0; p = 0.594; cl = -0.01; cu = 0.01 | d = -0.01; p = 0.26; cl = -0.01; cu = 0 |
| V. ord (FS144) | d = 0.01; p = 0.27; cl = -0.01; cu = 0.02 | d = 0; p = 0.94; cl = -0.02; cu = 0.02 | d = -0.01; p = 0.071; cl = -0.03; cu = 0 | d = 0; p = 0.693; cl = -0.02; cu = 0.01 | d = -0.01; p = 0.423; cl = -0.02; cu = 0.01 | d = 0; p = 0.981; cl = -0.02; cu = 0.02 |
| V.nat (14048) | d = 0.03; p = 0.006; cl = 0.01; cu = 0.05 | d = 0.03; p = 0.007; cl = 0.01; cu = 0.06 | d = 0; p = 0.902; cl = -0.02; cu = 0.02 | d = 0.03; p = 0.038; cl = 0; cu = 0.05 | d = 0.04; p = 0.005; cl = 0.01; cu = 0.07 | d = 0.02; p = 0.125; cl = -0.01; cu = 0.04 |
| V. kan (5S149) | d = -0.01; p = 0.365; cl = -0.03; cu = 0.01 | d = -0.02; p = 0.086; cl = -0.04; cu = 0 | d = -0.02; p = 0.018; cl = -0.04; cu = 0 | d = -0.01; p = 0.329; cl = -0.03; cu = 0.01 | d = -0.01; p = 0.455; cl = -0.03; cu = 0.01 | d = 0; p = 0.967; cl = -0.02; cu = 0.02 |
| V. len (5F79) | d = -0.01; p = 0.582; cl = -0.05; cu = 0.03 | d = -0.02; p = 0.18; cl = -0.06; cu = 0.01 | d = -0.01; p = 0.708; cl = -0.04; cu = 0.03 | d = -0.01; p = 0.439; cl = -0.05; cu = 0.02 | d = -0.01; p = 0.639; cl = -0.05; cu = 0.03 | d = -0.01; p = 0.665; cl = -0.04; cu = 0.03 |
| V. rum (1S45) | d = -0.04; p = 0.544; cl = -0.18; cu = 0.09 | d = -0.06; p = 0.4; cl = -0.19; cu = 0.08 | d = -0.05; p = 0.481; cl = -0.18; cu = 0.09 | d = -0.14; p = 0.039; cl = -0.28; cu = -0.01 | d = -0.1; p = 0.165; cl = -0.25; cu = 0.04 | d = -0.12; p = 0.073; cl = -0.26; cu = 0.01 |
| V. nor (1F211) | d = -0.08; p = 0.103; cl = -0.18; cu = 0.02 | d = -0.04; p = 0.46; cl = -0.13; cu = 0.06 | d = -0.12; p = 0.022; cl = -0.21; cu = -0.02 | d = 0; p = 0.938; cl = -0.1; cu = 0.09 | d = -0.03; p = 0.625; cl = -0.14; cu = 0.08 | d = 0; p = 0.973; cl = -0.1; cu = 0.1 |
| V. spl (13B01) | d = 0.03; p = 0.043; cl = 0; cu = 0.06 | d = 0.02; p = 0.126; cl = -0.01; cu = 0.05 | d = -0.03; p = 0.054; cl = -0.05; cu = 0 | d = -0.02; p = 0.2; cl = -0.05; cu = 0.01 | d = -0.03; p = 0.087; cl = -0.06; cu = 0 | d = -0.01; p = 0.348; cl = -0.04; cu = 0.01 |
| V. spl (1A01) | d = 0.02; p = 0.002; cl = 0.01; cu = 0.04 | d = 0.07; p = 0; cl = 0.05; cu = 0.09 | d = 0; p = 0.486; cl = -0.01; cu = 0.02 | | | |
| V. tas (1F187) | d = 0.01; p = 0.698; cl = -0.05; cu = 0.07 | d = 0; p = 0.823; cl = -0.06; cu = 0.06 | | | | |

**Table S5: ANOVA results endochitinase (related to Figure S5)**

ANOVA table of differences in endochitinase activity for combinations of various degraders (y-axis) in co-culture with different cross-feeders (x-axis). Posthoc test for multiple comparisons of means using Fisher LSD of a two-way ANOVA (with the factors culture-type and day of the experiment) revealed these scores. Shown are calculated differences in endochitinase activity mono- and co-culture (d), the associated *p*-value (p), and the lower and upper 95% confidence intervals (cl, cu). In cases where the rounded digits indicate p = 0, p < 0.0005. In cases where the rounded digits indicate d = 0, d < 0.005.

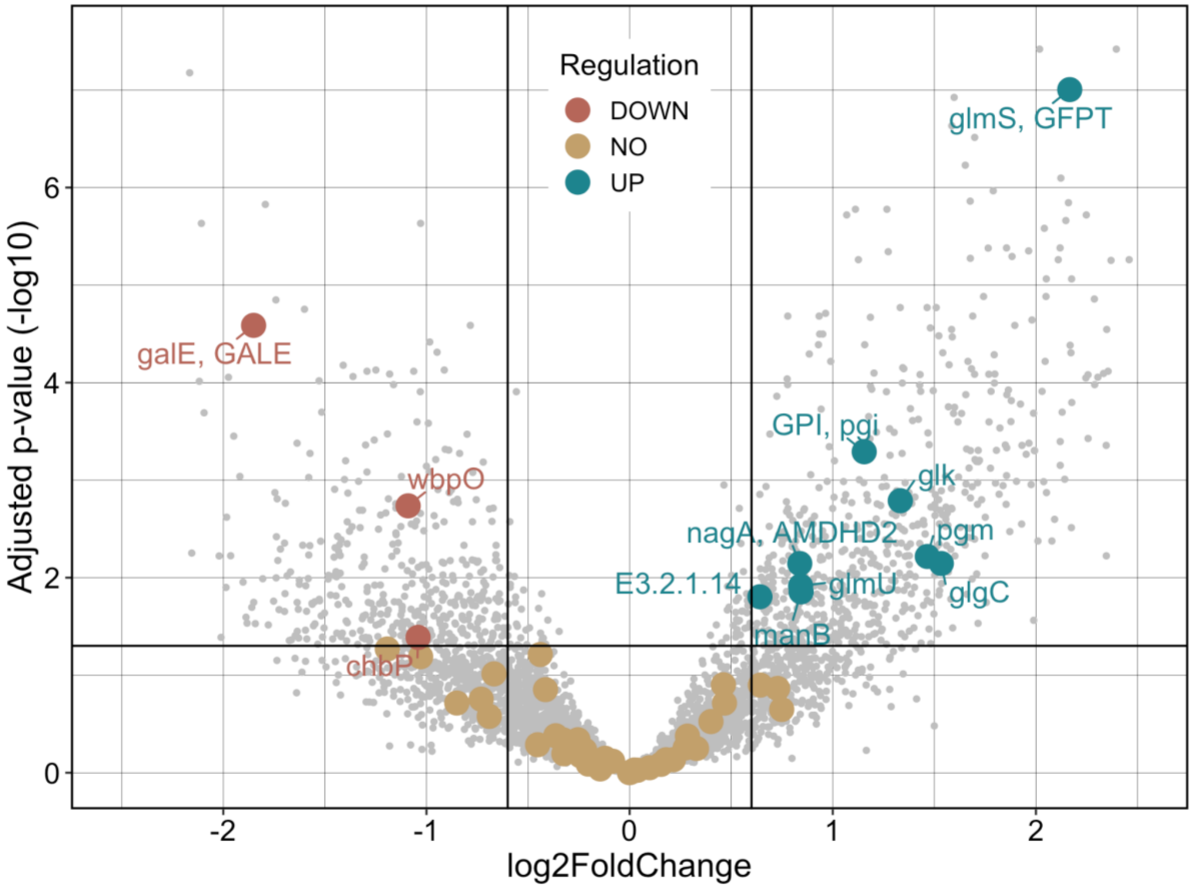


**Figure S6: *Alteromonas* induces chitin metabolism pathway**

Differences in amino sugar and nucleotide sugar pathway expression of *V. natriegens* in the presence of *Alteromonas*. Dots indicate individual genes of that pathway found in the genome of *V. natriegens*, black lines indicate significance cut offs (vertical lines indicate +/- 0.6 log2 Fold Change, the horizontal line indicates $p < 0.05$) colours indicate differential expression based on the cut off criteria; down-regulation (red), no difference (yellow), up-regulation (green). In total eleven genes were differentially expressed in the presence of *Alteromonas* compared to a degrader monoculture. Two were down regulated while nine were upregulated. This indicates that not only enzymes but the complete pathway of chitin degradation and consumption is stimulated by the presence of *Alteromonas*.

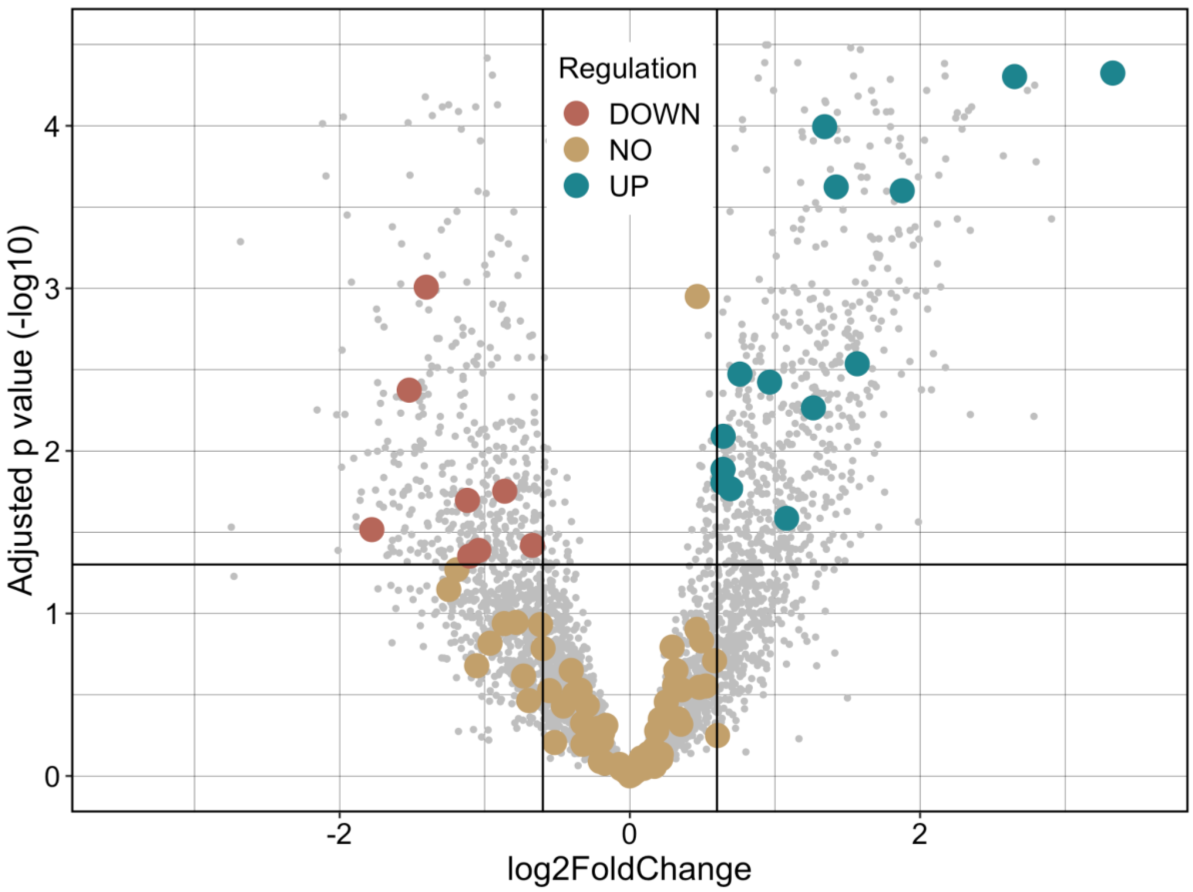


**Figure S7: *Alteromonas* influences expression of glycosyl hydrolases in *V. natriegens*.** Differences in glycosyl hydrolase expression in *V. natriegens* in the presence vs absence of *Alteromonas*. Dots indicate individual glycosyl hydrolases found in the genome of *V. natriegens*, black lines indicate significance cut offs (vertical lines indicate |log2 Fold Change| = 0.6, the horizontal line indicates $p < 0.05$) colors indicate differential expression based on the cut off criteria; down-regulation (red), no difference (yellow), up-regulation (green). 14 out of 87 glycosyl hydrolase were significantly upregulated in the presence of *Alteromonas*. Eight out of 87 glycosyl hydrolases were significantly down-regulated in the co-culture compared to the degrader monoculture.

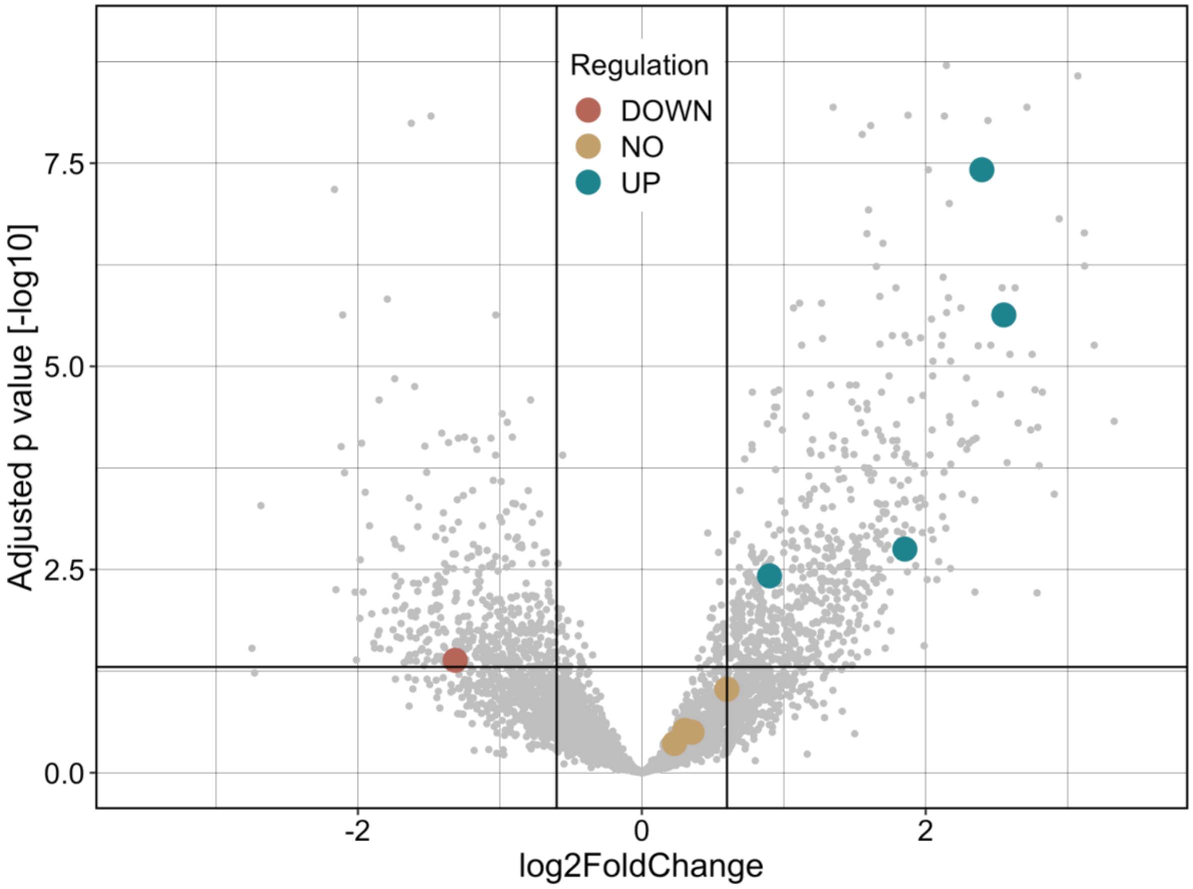


**Figure S8: *Alteromonas* induces expression of auxiliary enzymes in *V. natriegens.***

Differences in the expression of auxiliary enzymes in *V. natriegens* in the presence vs absence of *Alteromonas*. Dots indicate individual auxiliary enzymes found in the genome of *V. natriegens*, black lines indicate significance cut offs (vertical lines indicate |log2 Fold Change| = 0.6, the horizontal line indicates $p < 0.05$) colors indicate differential expression based on the cut off criteria; down-regulation (red), no difference (yellow), up-regulation (green). Four out of nine auxiliary enzymes were significantly upregulated in the presence of *Alteromonas*. One out of nine auxiliary enzymes were significantly down-regulated in the co-culture compared to the degrader monoculture.

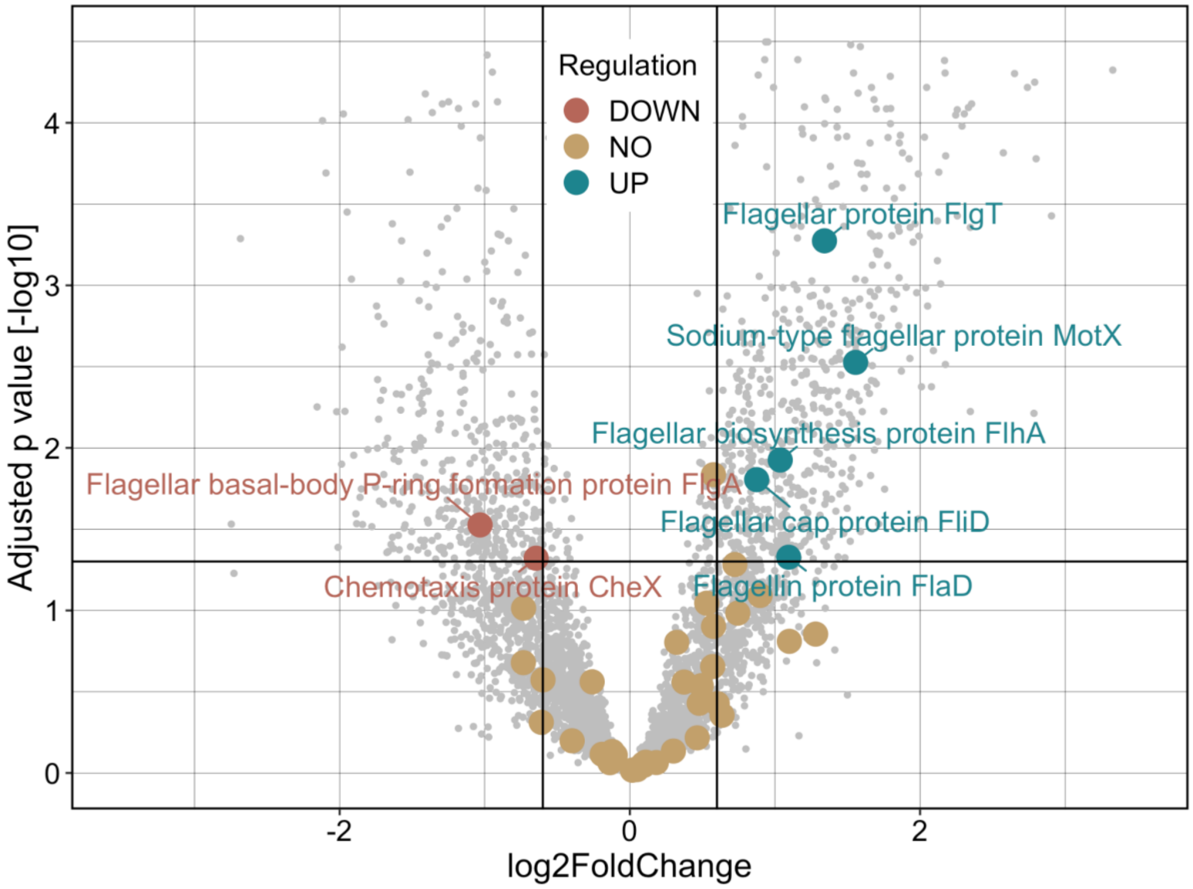


**Figure S9: *Alteromonas* slightly increases expression of motility gene cluster in *V. natriegens*.** Differences in motility gene expression in *V. natriegens* in the presence vs absence of *Alteromonas*. Dots indicate individual motility genes found in the genome of *V. natriegens* (Methods), black lines indicate significance cut offs (vertical lines indicate |log2 Fold Change| = 0.6, the horizontal line indicates $p < 0.05$) colors indicate differential expression based on the cut off criteria; down-regulation (red), no difference (yellow), up-regulation (green). Five out of 48 motility genes were significantly upregulated in the presence of *Alteromonas*. Two out of 48 motility genes were significantly downregulated.

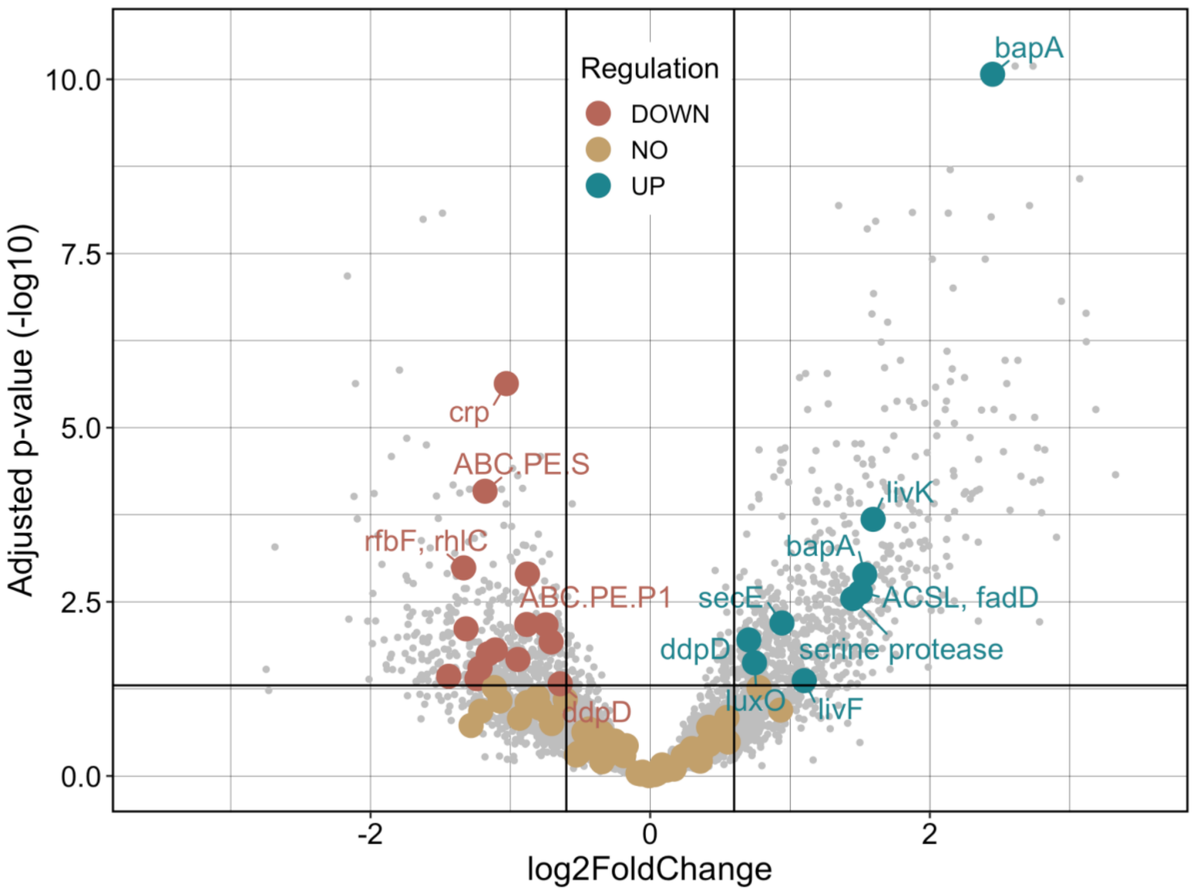


**Figure S10: *Alteromonas* decreases expression of quorum sensing genes in *V. natriegens*.** Differences in quorum sensing gene expression in *V. natriegens* in the presence vs absence of *Alteromonas*. Dots indicate individual quorum sensing found in the genome of *V. natriegens* (Methods), black lines indicate significance cut offs (vertical lines indicate |log2 Fold Change| = 0.6, the horizontal line indicates $p < 0.05$) colors indicate differential expression based on the cut off criteria; down-regulation (red), no difference (yellow), up-regulation (green). Nine out of 282 quorum sensing genes were significantly upregulated in the presence of *Alteromonas*. Fifteen out of 282 quorum sensing genes were significantly downregulated.

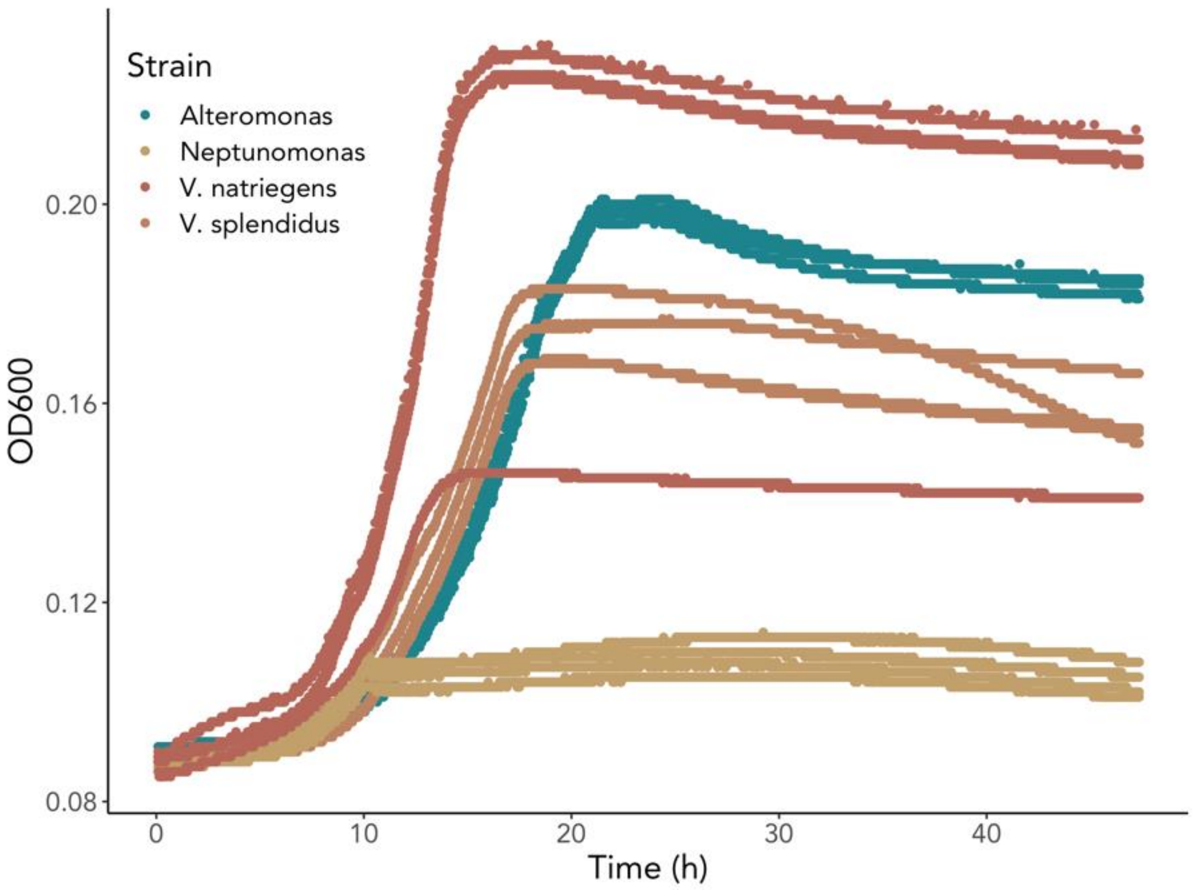


**Figure S11: Growth of degraders and cross-feeders on chitinase.**

Microbial species are able to utilize enzymes as carbon sources. *Alteromonas* (green), *Neptunomonas* (yellow), *V. natriegens* (red), *V. splendidus* (brown) display differing growth dynamics on chitinases as the sole carbon source.

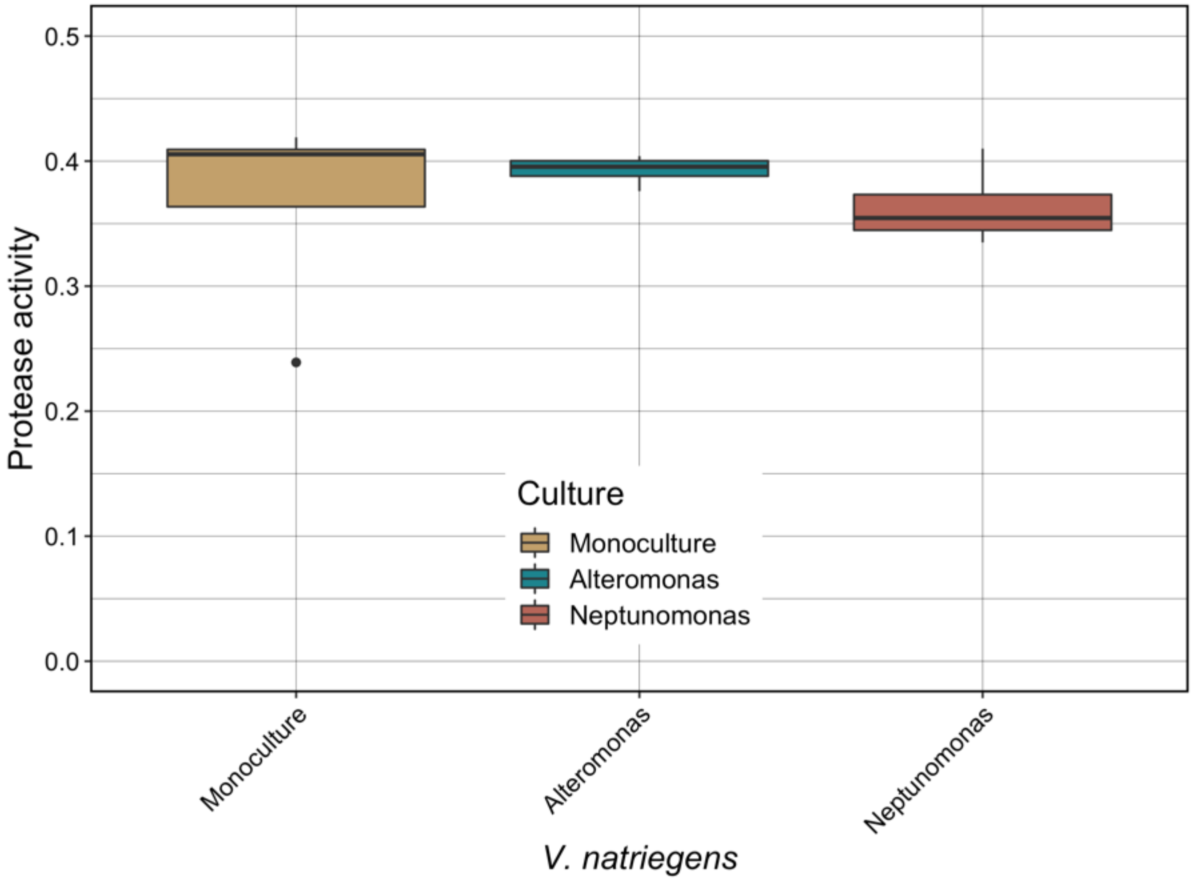


**Figure S12: Communities secrete proteases.**

Protease activity of *V. natriegens* monocultures (yellow) compared to co-culture with *Alteromonas* (green) and *Neptunomonas* (red). When grown on chitinase as a sole carbon source, communities will secrete proteases.

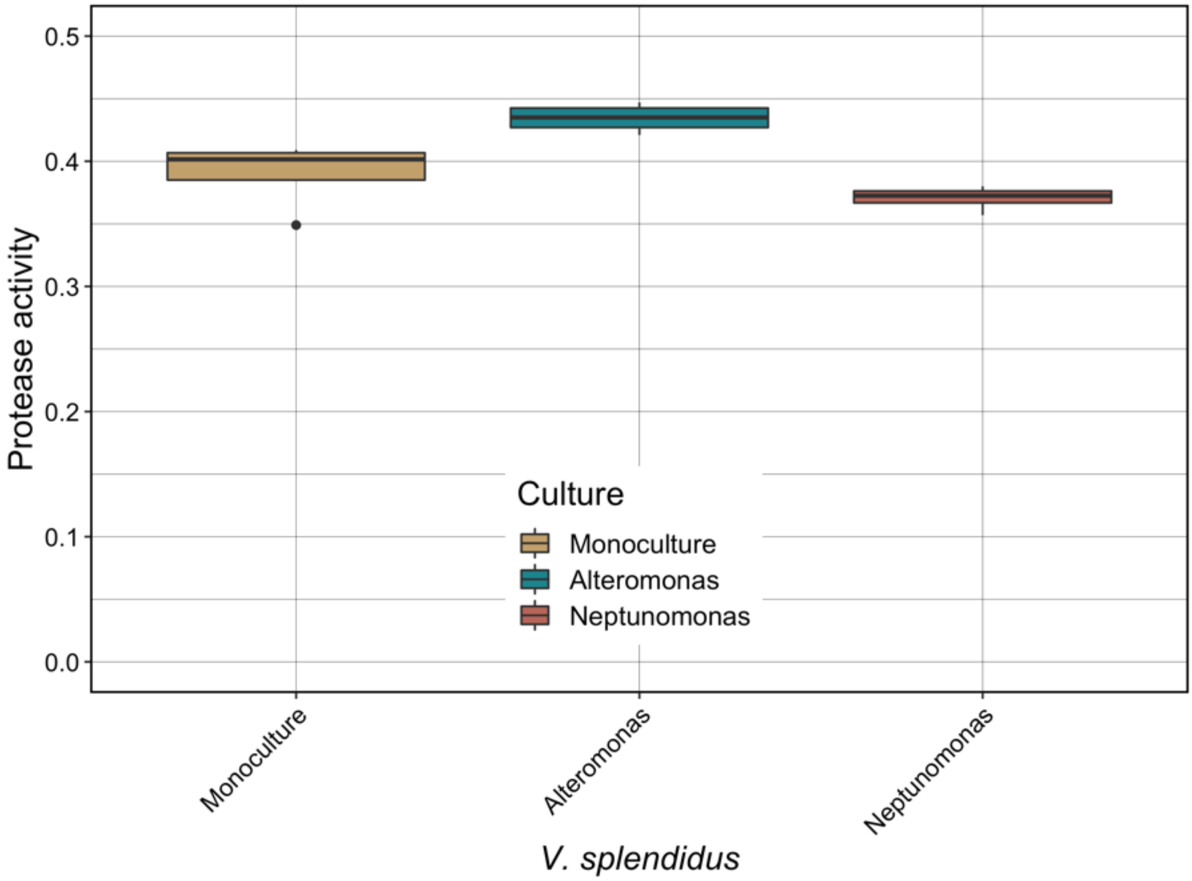


**Figure 13: Communities secrete proteases.**

Protease activity of *V. splendidus* monocultures (yellow) compared to co-culture with *Alteromonas* (green) and *Neptunomonas* (red). When grown on chitinase as a sole carbon source, communities will secrete proteases.
